# Evolutionary rate covariation across malaria parasite species enables inference of protein interactions

**DOI:** 10.64898/2026.01.29.702598

**Authors:** Helena D Hopson, Radoslaw Igor Omelianczyk, Anayansi Ramirez, Jordan H. Little, Nathan Clark, Paul A Sigala, Ellen M Leffler

**Author notes:** Corresponding authors: Ellen M. Leffler, Helena D. Hopson.

## Abstract

Despite publication of the *Plasmodium falciparum* reference genome over 20 years ago, one-third of its genes remain functionally unannotated, and information is limited for many others. Proteins that act in the same pathway or complex tend to experience similar shifts in evolutionary pressure, such that correlated constraints across species can indicate co-functional proteins. To investigate this connection, we calculated relative evolutionary rates across the genome on a phylogeny of 22 *Plasmodium* species and assigned each protein-protein pair a score representing the strength of evolutionary rate covariation (ERC). We show that known pathways and interacting proteins across lifecycle stages in *Plasmodium* have strong ERC signals. By scanning genome-wide for additional proteins showing high ERC with established interacting proteins, we find enrichment of stage expression and physically interacting protein pairs supporting new candidate functions for proteins. More generally, we demonstrate the utility of ERC to prioritize proteins for hypothesis-driven functional follow up by showing that a protein with little functional characterization (PF3D7_0811600) shows high ERC and co-localizes with high molecular weight rhoptry proteins 2 and 3 (RhopH2 and RhopH3), which form an ion channel that enhances parasite permeability of infected red blood cells. The ERC matrix can be queried to extract *Plasmodium* proteins showing high ERC with any protein of interest to prioritize candidate genes and accelerate discovery of novel functional connections.

## Introduction

*Plasmodium* was responsible for 263 million malaria cases and 597,000 deaths in 2023, the majority of which were caused by *Plasmodium falciparum* [1]. Despite decades of research, around one-third of *P. falciparum* genes lack functional annotation [2], in part due to limited tools for genetic manipulation and evolutionary distance from common model organisms [3]. Recently, high-throughput mutagenesis screens have been performed in *P. falciparum* [4], the rodent parasite *P. berghei* [5], and the zoonotic monkey parasite *P. knowlesi* [6, 7], resulting in gene-level assignments of essentiality and impact on growth. Additionally, a protein-protein interaction network in *Plasmodium* derived from co-migration of protein complexes in native gels queried physical protein-protein interactions among about half of *Plasmodium*’s ∼5,400 proteins [8, 9]. A major goal of *Plasmodium* molecular research is to ascribe more detailed functions to proteins and new functions to uncharacterized genes toward identifying future drug and vaccine targets. Here, we aim to complement existing experimental screens with an evolutionary approach to identify functionally related proteins.

Over the course of evolution, species experience distinct environments and evolutionary pressures. The extent of purifying selection can be quantified as an evolutionary rate, measured as the number of amino acid substitutions per unit time along a branch. A relative evolutionary rate (RER) represents the branch-specific evolutionary rate for a particular protein relative to its average rate of evolution across all branches of a species tree [10, 11]. Additionally, the gene-branch rate is normalized by the average rate on that branch across all genes. In its final form, an RER represents how much faster or slower a protein is evolving in one species relative to all species in the phylogeny. Proteins that have similar functions or act in the same pathway tend to experience similar shifts in evolutionary pressures over time [12-14], such that their rates tend to increase (or decrease) on the same branches of the phylogeny. Evolutionary rate covariation (ERC) quantifies the correlation of two proteins’ RERs across a species tree and can thus be used to identify co-functional proteins [12, 15]. A high ERC value between a pair of proteins can be driven by direct (i.e. physical) or indirect (i.e. shared pathway or function) protein interactions [12, 16]. This ERC-driven approach to identify co-functional proteins has been applied in a wide range of taxa including mammals [17-19], *Drosophila* [20-22], and yeast [16, 23]. Several of these studies have shown experimental validation of ERC-predicted candidates such as connecting new proteins to pre-defined pathways [17, 20, 22] and localization of an uncharacterized protein [19]. Furthermore, in mammals, ERC performed just as well if not better than functional screens [17].

*Plasmodium* species have faced strong and variable evolutionary pressures due to their parasitic lifestyle and frequent host switches, making evolutionary constraint a potentially powerful approach to help predict protein function in these taxa. The *Plasmodium* phylogeny includes parasite species that infect all classes of terrestrial vertebrates and a range of mosquito vectors, with varying degrees of host specificity. The recent increase in the number of reference genomes available for divergent *Plasmodium* species including avian, ape, monkey, and rodent infecting species now make ERC possible in this clade. Here, we calculate ERC for 4,360 proteins and use it to infer new protein functions in *Plasmodium*. We show that there is indeed strong ERC signal among known pathways and co-functional proteins, and that scanning genome-wide reveals clusters of functionally related proteins in networks of high ERC, supported by enrichment in functional datasets and published experimental data. Furthermore, we use ERC to identify a link between a protein with little functional characterization (PF3D7_0811600) and components of a parasite permeability pathway and show that these proteins co-localize. This connection illustrates the utility of ERC as a screening tool to prioritize protein candidates for functional follow up. We establish ERC as an accessible resource to query proteins and extract high ERC pairs through an Rshiny app (https://lefflerlab.chpc.utah.edu/erc_plasmodium/). Notably, proteins of all 22 *Plasmodium* species included in this study can be queried for ERC interactions. This resource will accelerate the identification of functional roles for uncharacterized proteins and co-functional relationships between proteins, facilitating a more comprehensive understanding of *Plasmodium* biology and discovery of new antimalarial drug and vaccine targets.

## Results

### An evolutionary rate covariation resource for *Plasmodium*

To calculate evolutionary rate covariation, amino acid sequences of 22 *Plasmodium* species (*P. adleri, P. reichenowi, P. praefalciparum, P. relictum, P. vinckei vinckei vinckei, P. vivax-like, P. vivax, P. yoelii yoelii, P. ovale curtisi, P. malariae, P. knowlesi, P. inui, P. gallinaceum, P. gaboni, P. fragile, P. berghei, P. billcollinsi, P. blacklocki, P. chabaudi chabaudi, P. coatneyi, P. cynomolgi*, and *P. falciparum)* were downloaded from PlasmoDB [24] (Table S1). Proteins were assigned to 8,085 ortholog groups using OrthoMCL, and groups with at least 10 1-to-1 orthologs were retained. Following steps used to generate ERC matrices in other taxa [22], protein sequences of 1-to-1 orthologs in each group (excluding paralogs) were aligned and RERs were estimated across the phylogeny (Fig 1A). ERC between each pair of ortholog groups with at least 10 shared species was estimated as the pairwise correlation between their RERs (Fig 1B). Because 1-to-1 orthology is required for rate estimation, large, duplicated gene families (such as var genes) are absent from this dataset. However, we retain genes with lineage-specific duplications that have sufficient 1-to-1 orthology elsewhere in the phylogeny.

**Fig 1.**
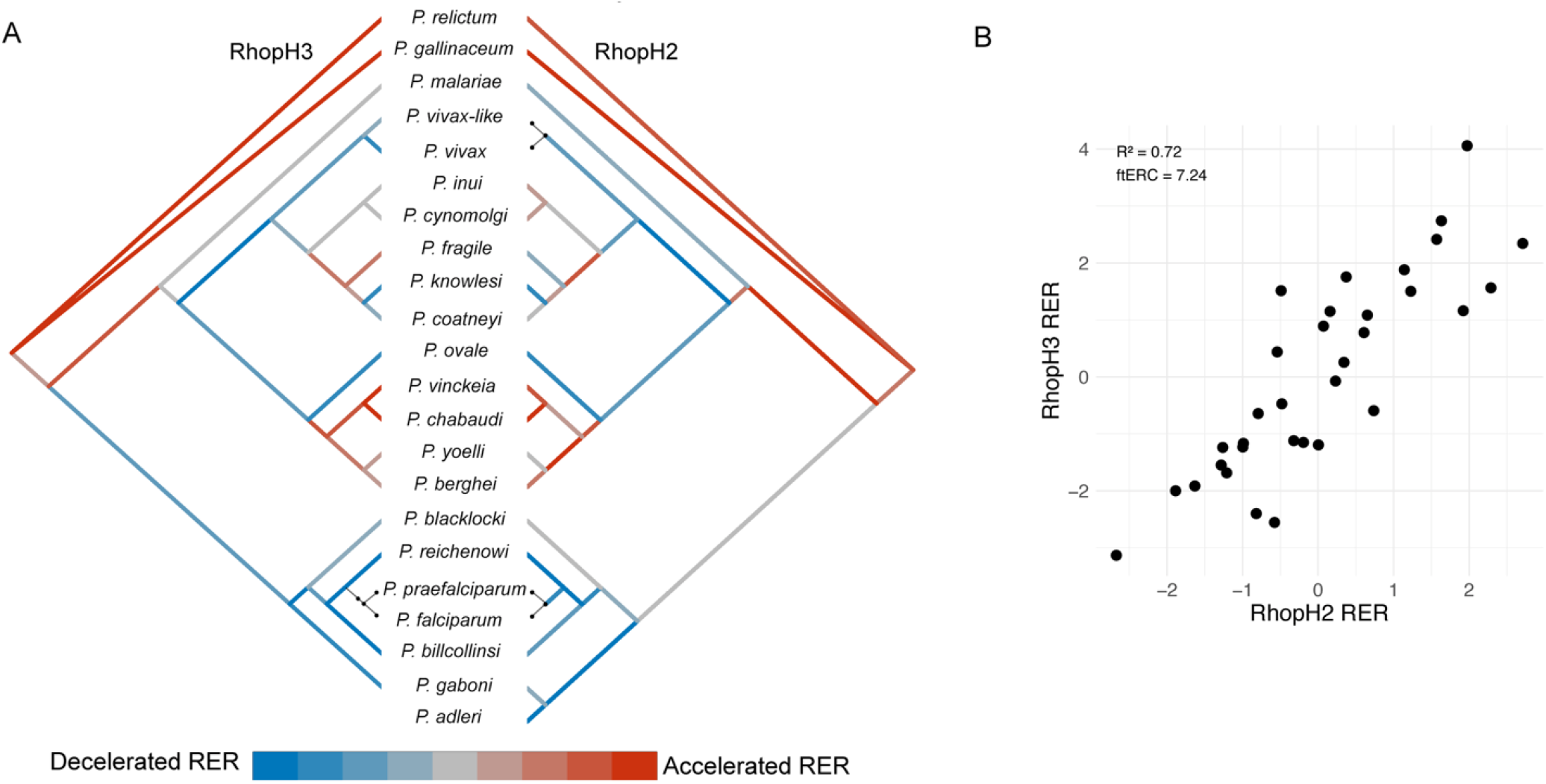
Phylogeny and example of high ERC between RhopH2 and RhopH3. (A) The species topology of the 22 *Plasmodium* species used to create the ERC matrix. Dots connect species names to branches. Branches are colored by relative evolutionary rate (RER) for RhopH3 (left) and RhopH2 (right), with red indicating higher rates (less constraint) and blue indicating lower rates, (more constraint). Terminal branches with thin black lines indicate that there were too few substitutions to calculate RER. RhopH3 is missing a terminal branch to *P. gaboni* because there was not a 1:1 ortholog assigned. (B) Scatter plot of RER values for RhopH2 and RhopH3 with the Pearson’s r correlation coefficient and fisher transformed ERC value.

The resulting matrix contains 9,370,728 ERC values across 4,360 proteins. Fisher transformed ERC values ranged from -7.2 to 8.9 and the median and mean values were close to 0 (Fig S1). In total, 4,134 of the ortholog groups (94%) had a single-copy ortholog in *P. falciparum*. Hereafter, we refer to genes by *P. falciparum* gene name and by another representative species for those without single-copy *P. falciparum* orthologs. However, because at least 10 species are represented in each ortholog group, ERC captures proteins that are functionally related across multiple *Plasmodium* species.

ERC is a rank-based approach to prioritize co-functional interactions, such that the higher the ERC value the more likely two proteins functionally interact. Significance is assessed by comparison to the distribution of all ERC values genome-wide. For example, we observed ERC of 7.2, in the 99.9th percentile of all genome-wide ERC values, between RhopH2 and RhopH3, which form an established complex (Fig 1B) [25, 26]. To assess significance of pathways with multiple proteins, we compared the mean of all pairwise ERC values to those of 1000 random protein sets of the same size.

### Elevated ERC in known *Plasmodium* pathways

We selected six pathways that are either well-studied in *Plasmodium* (fatty acid synthesis, hemoglobin digestion, red blood cell (RBC) invasion) or conserved across eukaryotes and have previously shown strong ERC in other taxa (DNA replication, homologous recombination, mismatch repair) [17, 23] (Table S2). We found elevated ERC among proteins involved in five of these pathways (Table 1, empirical p < 0.018, Fig S2-5) indicating that, as in other taxa, many pathways have a significant ERC signature in *Plasmodium*. We did not observe significantly higher ERC among proteins in the mismatch repair set (Table 1, empirical p = 0.5, Fig S6).

**Table 1.** Mean of all pairwise ERC values between proteins in *Plasmodium* pathways.

| Pathway | Mean ERC | Empirical p-value | Number of genes in ERC matrix |
| --- | --- | --- | --- |
| **DNA replication** | 0.37 | < 0.001 | 41 |
| **Homologous recombination** | 0.58 | 0.012 | 12 |
| **Fatty acid synthesis** | 0.27 | 0.017 | 25 |
| **Hemoglobin digestion** | 0.53 | 0.002 | 19 |
| **Mismatch repair** | 0.07 | 0.462 | 28 |
| **RBC invasion** | 0.95 | < 0.001 | 77 |

To assess ERC signal in other groups of functionally related proteins, we tested for significant mean pairwise ERC among STRING network clusters containing 3-500 proteins [27], filtering to keep only proteins also in the ERC dataset. Of these clusters, 34% (262/765) had significant mean pairwise ERC (empirical p < 0.05) (Table S3). In addition, the mean pairwise ERC of 17,174 protein-protein interactions predicted in a *Plasmodium* interactome based on experimental co-migration was significant (0.34, empirical p < 0.001) [8]. Taken together, these results indicate that functionally related proteins and pathways in *Plasmodium* show significant ERC. Thus, these proteins and pathways represent good candidates for an ERC driven approach for prioritization of unannotated functional connections, as subsequently described.

### Genome-wide scan for additional contributors to red blood cell invasion

For pathways where components have elevated ERC, the entire ERC dataset can then be searched for additional proteins showing similarly high ERC with proteins in the pathway, as a screen for a new set of candidates that may perform an unrecognized function in the same pathway. We applied this ERC-guided approach to uncover new roles of proteins in red blood cell (RBC) invasion. RBC invasion is essential for parasite growth and both an antimalarial drug and vaccine target, yet many proteins and complex components remain undiscovered [28-31]. To this end, we compiled a list of 131 proteins that have been linked to RBC invasion based on functional or localization information from review papers and the Malaria Parasite Metabolic Pathway (MPMP) database [28, 32-34] (Table S2). Of the curated RBC invasion protein list, 77 were present in the ERC matrix, spanning multiple stages from pre-to post-invasion, including surface, microneme, and rhoptry proteins. RBC invasion proteins absent from the ERC matrix include those in multigene families without sufficient 1-to-1 orthology for this approach, such as stevor, SERA, and CLAG genes.

The mean pairwise ERC of the 77 RBC invasion proteins was significantly elevated (0.95, empirical p < 0.001) (Table 1, Figure S7). ERC was especially high among a subset of 19 RBC invasion proteins (Fig 2A). Within this subset, most RBC invasion proteins had their highest genome-wide ERC values with other RBC invasion proteins. For example, RBC invasion protein RhopH3 had six out of its 20 highest ERC values with other RBC invasion proteins in the curated list, including RhopH2 (Fig 2B).

**Fig 2.**
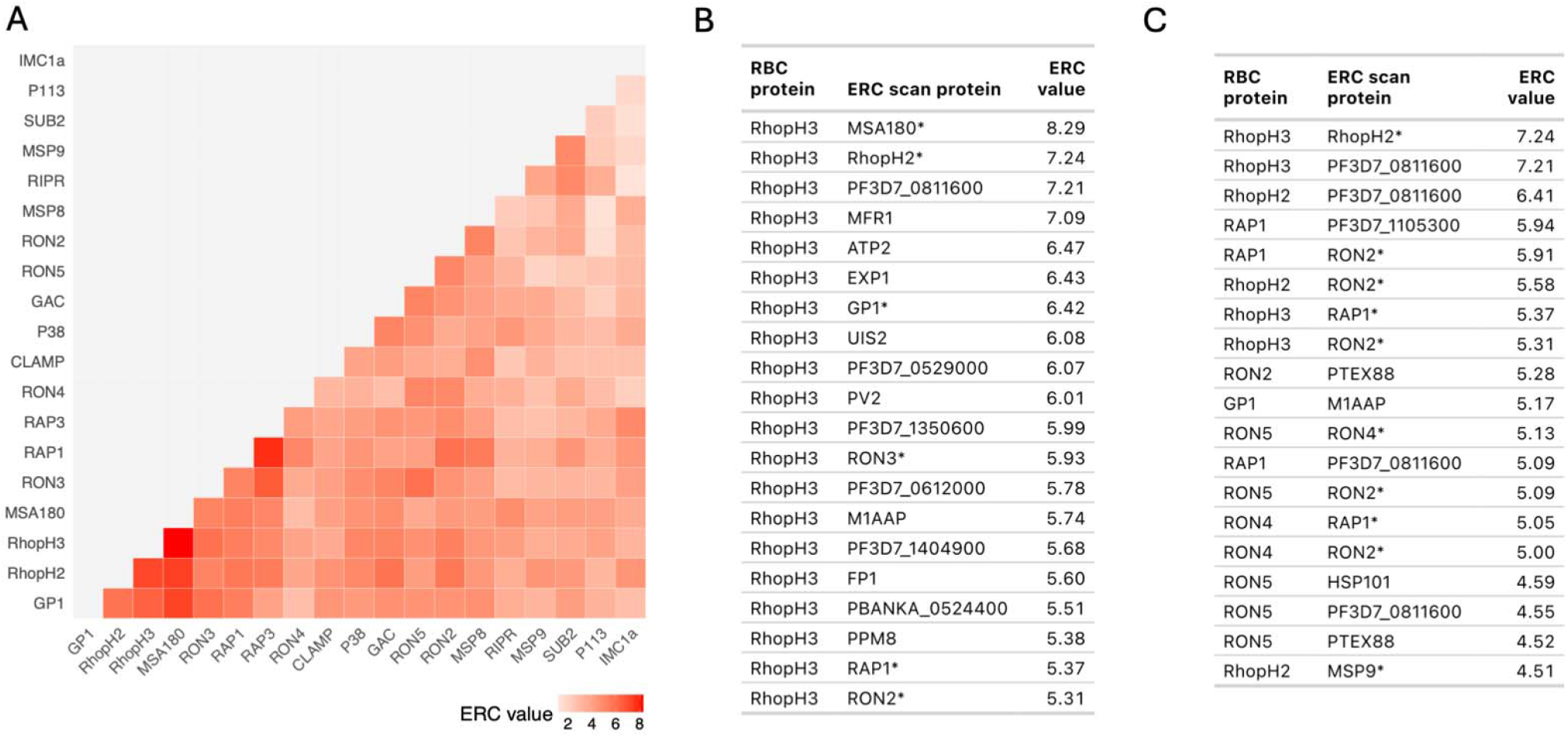
Elevated ERC in RBC invasion proteins. (A) Heatmap showing high ERC in a subset of 19 RBC invasion proteins (full heatmap in Fig S7). (B) Table of the 20 proteins showing the highest ERC with RhopH3 genome wide. (C) Table of the 19 pairs from the RBC invasion – genome ERC scan that also pair in the interactome. Proteins in the curated RBC invasion list are marked with an asterisk. All ERC values in the tables are in the 99.9th percentile of all ERC values genome wide.

To scan for proteins with potential functions in RBC invasion, we took proteins with an ERC value in the 99.9th percentile of all values (ERC > 4.48) with any RBC invasion protein, resulting in 550 RBC invasion - genome protein pairs. Of those pairs, 66 had two RBC invasion proteins (12%) and 484 had one protein not in the RBC invasion list, for a total of 228 unique proteins without a described role in RBC invasion (Table S4). These 228 new candidates for RBC invasion were enriched for expression in asexual blood stages compared to all ERC proteins (Fig S8). To compare our results with previous scans for RBC invasion proteins, we intersected our candidates with those from a study which used an approach based on transcriptional similarity to identify a network of 418 candidate RBC invasion proteins (387 after converting to newer gene names and excluding reference genes, 303 in ERC matrix) [35]. Of our 228 RBC invasion candidates, 39 were also identified in this co-expression study (Table S5). Only seven of the 26 reference RBC invasion genes this study used to identify co-expressed genes were present in the ERC matrix. Scanning for proteins in the 99.9th ERC percentile with these seven genes resulted in 98 candidates, of which 20 (20%) were also identified in the co-expression study. Next, we sought to compare our protein pairs with an experimentally derived *Plasmodium* protein-protein interactome [8]. Of the 550 RBC invasion - genome scan protein pairs, 368 contained proteins that were both included in the protein-protein interactome study. Of these, 19 pairs also interacted in the interactome study (Fig 2C). Out of 1,000 permutations of random sets of 368 ERC protein-protein pairs, a number as high as 19 shared protein pairs between ERC and interactome was never observed (empirical p<0.001). This indicates that high ERC with RBC invasion proteins is enriched for physical interactions. Together, these results suggest ERC is identifying both direct (e.g. physical) and indirect (e.g. co-expression) interactions as well as both known and novel interactions, the latter indicating candidate proteins newly implicated in RBC invasion.

### Genome-wide ERC clustering

We identified groups of proteins linked by high ERC by clustering based on the 99.9^th^ percentile of ERC values (> 4.48), consisting of 18,742 protein pairs (Table S6). The clustering resulted in 360 clusters with at least three proteins, covering about one-third of ortholog groups in the ERC matrix (1,474/4,360) (Table S7). To begin to link ERC clusters with potential functions, we tested for functional enrichment using STRING database annotations (including Gene Ontology, KEGG, Reactome, UniProt, STRING Network Clusters) and Gene Ontology annotations accessed via the org.Pf.plasmo.db Bioconductor package. Functional enrichment analysis of the 326 clusters with at least three *P. falciparum* proteins revealed 47 clusters with at least one enriched functional term (Fig 3, Table S8, S9). Additionally, 59 pairs of proteins within ERC clusters had a confident STRING interaction (combined score > 400 and experimental evidence score > 0) (Fig 3, Table S10).

**Fig 3.**
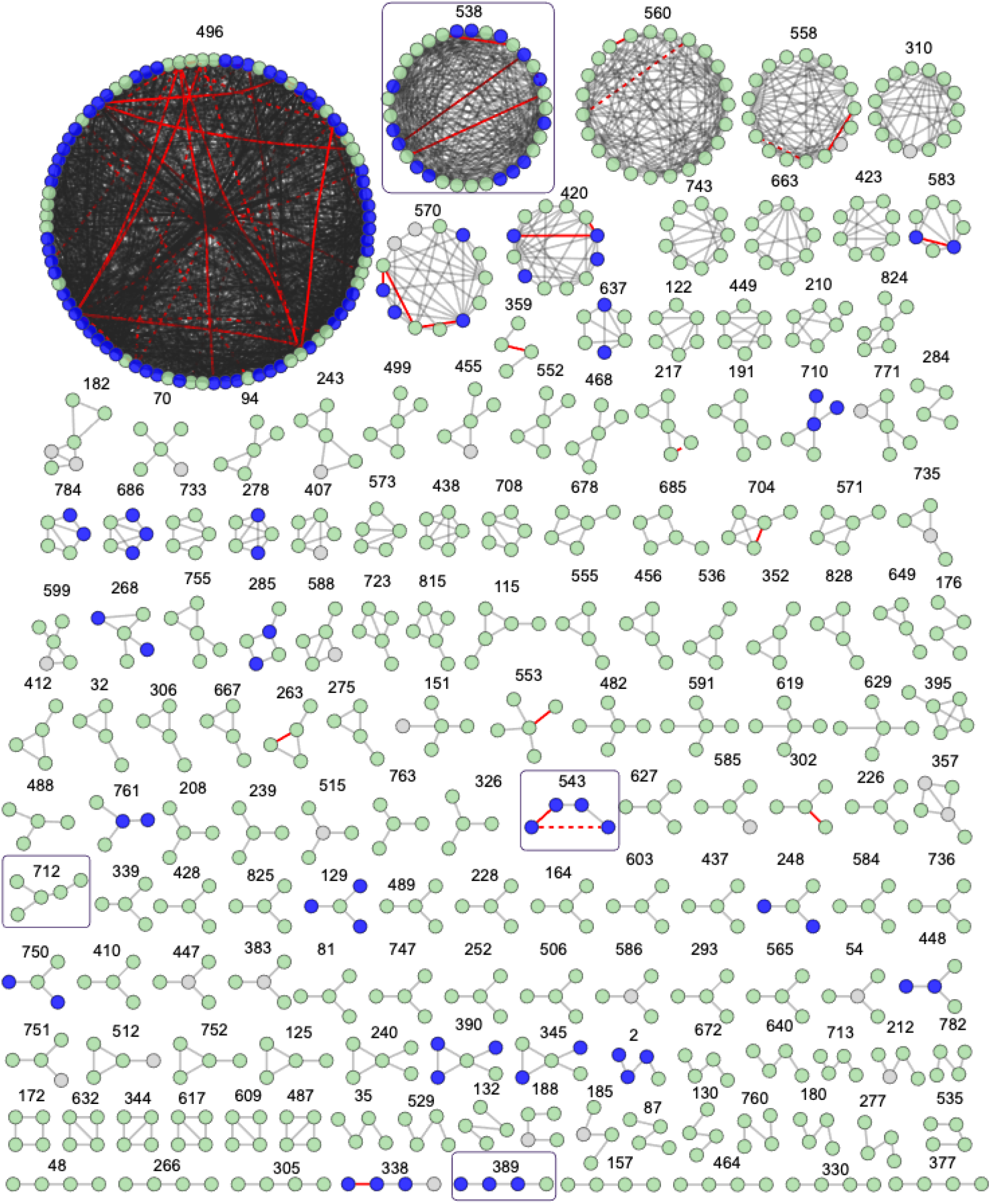
Clustering of the highest ERC genome wide. Clustering of the 99.9th percentile of all ERC values (ERC > 4.48). Only clusters with at least 4 proteins are shown. Grey edges indicate that the ERC value between proteins is above this value. Red edges indicate a confident interaction reported in STRING (combined score >400 and experimental evidence > 0), solid indicates both a STRING interaction and ERC > 4.48 and dashed indicates a STRING interaction without an ERC value > 4.48. Purple nodes indicate a *P. falciparum* ortholog that is part of an enriched functional term within the cluster (Table S8, S9) and white nodes represent an ortholog group without a *P. falciparum* ortholog. Clusters shown in subsequent figures are boxed.

To assess if the clusters indicate new connections between unannotated proteins and proteins with established functions, we searched for clusters containing proteins within the pathways from Table 1. Cluster 633 includes two proteins in the hemoglobin digestion pathway (Table S2), DPAP1 and APP, along with a third protein, PF3D7_0104200, which is a lipid transporter and does not have a known role in hemoglobin digestion (Fig 4A). Lipids are thought to be involved in detoxification of by-products from hemoglobin digestion [36], and both PF3D7_0104200 and DPAP1 localize within the parasite [37], supporting a potential role in hemoglobin digestion (Fig 4A). Cluster 543 includes three proteins included in the Table 1 pathways, including DNA replication and homologous recombination (Table S2). All four proteins in the cluster are annotated with the GO term helicase activity (Fig 4B, Table S8). Three of the four proteins are annotated as “putative”, indicating that their annotations are likely inferred from sequence homology to proteins in other species and have not been experimentally validated. Accordingly, we unable to find any publications on the functional role of these three proteins in *Plasmodium* (FANCJ/PF3D7_1408400/putative FANCJ-like helicase, POLQ/PF3D7_1331100/putative DNA polymerase theta, and PF3D7_1357500/putative DNA helicase). Correlated evolution of these “putative” proteins with experimentally validated *P. falciparum* helicase gene *RECQ1* across the *Plasmodium* phylogeny supports their functional conservation in *Plasmodium* as well.

**Fig 4.**
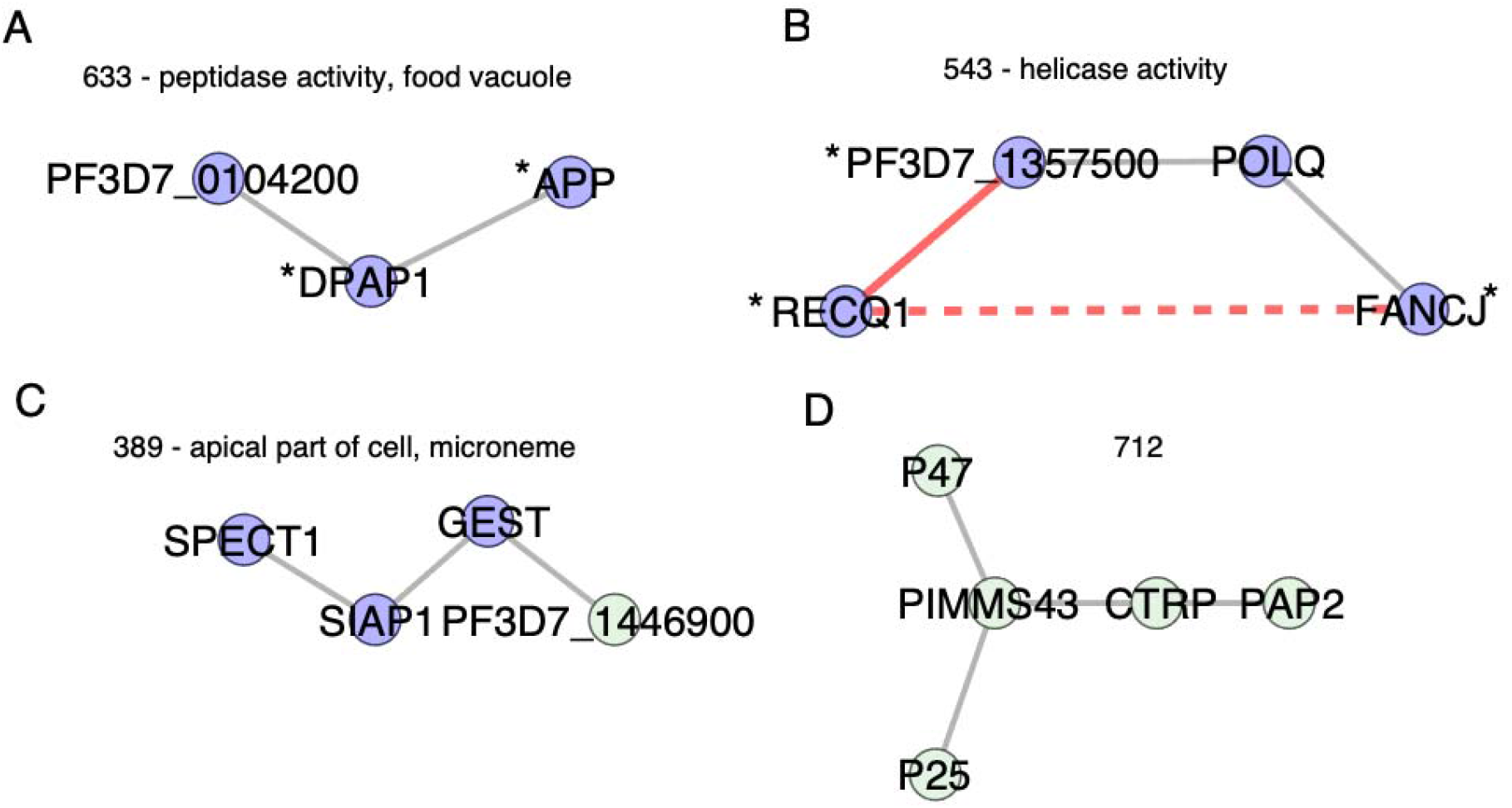
Examples of clusters containing functionally related proteins. Asterisk indicates proteins included in pathways tested in Table 1. Edges and nodes are formatted as described in Fig 3. (A) Cluster 633 has DPAP1 and APP from the hemoglobin digestion pathway and functional enrichment with GO term food vacuole and peptidase activity. (B) Cluster 543 has proteins included in DNA replication (RECQ1 and PF3D7_1357500) and homologous recombination (FANCJ) pathways from Table 1. All four proteins are annotated with GO annotation term helicase activity. (C) Cluster 389 has GO enrichment for microneme and apical part of cell. All four proteins have experimental evidence for functions in sporozoites and transmission from mosquito to liver stages. (D) Cluster 712 has no functional enrichment but all five proteins have experimental evidence for functions in ookinetes and invasion of mosquito midgut.

To assess if ERC based clustering identified co-functional genes specific to *Plasmodium* lifecycle stages outside the asexual blood stage, we looked for clusters with functional enrichment that contained proteins with functions in the mosquito. Cluster 389 contains three proteins annotated with GO term “apical part of cell” (SIAP1, SPECT1, GEST) and more specifically with “microneme” (Fig 4C, Table S8). All three proteins localize to the apical end of sporozoites, which mediate transmission from the mosquito to liver. SIAP1 has an early role in sporozoite migration to mosquito salivary glands, and SPECT1 and GEST are involved in later sporozoite traversal to vertebrate host liver cells [38-40]. The fourth protein in cluster 389 is an unnamed protein (PF3D7_1446900). It was previously identified as a putative glutaminyl-peptide cyclotransferase involved in evasion from the mosquito immune system by posttranslational modification of sporozoite surface proteins [41]. Selective pressures from the mosquito immune system could be the driving force of the correlated evolution in this cluster of co-functional proteins. Another example of a cluster of proteins that function in mosquito stages is cluster 712 (Fig 4D). This cluster does not have significant functional enrichment, but four of five proteins (P47, P25, PIMMS43, CTRP) have reported functions in mosquito immune evasion and are expressed on the surface of the ookinete, which invades mosquito epithelial cells [42-45]. P47 is essential for this process and is suggested to underly *Plasmodium* adaptation to new mosquito vector species, as specific combinations of parasite P47 haplotype and mosquito species determine successful infection of the mosquito [45]. High ERC within this cluster raises the possibility that the other proteins (P25, PIMMS43, CRTP) could be similarly involved in parasite adaptation to mosquitoes. Overall, these results highlight how ERC can identify co-functional proteins across different lifecycle stages and types of pathways, including those that are less accessible experimentally.

### ERC identifies new candidates interacting with RhopH2 and RhopH3

We identified five clusters in which at least one pair of proteins with high ERC were also found to interact in the experimentally derived interactome [8], for a total of 18 protein pairs (Table S11). Cluster 538 contains 12 such pairs and is enriched for GO terms including apical complex and rhoptry, which are the structures in the RBC-invading merozoite that facilitate initial binding and later discharge their contents into the host RBC during the invasion process [28]. This cluster includes six known rhoptry proteins: RAP1, RAP3, RON3, RON5, RhopH2 and RhopH3. RhopH2 and RhopH3 are part of the RhopH complex, a key component of the *Plasmodium* surface anion channel (PSAC) that is inserted on the infected host RBC membrane to facilitate nutrient uptake [29, 46, 47]. The scope of PSAC affiliated proteins is not known [46, 48], and this cluster suggests new proteins that could be involved. We focused on PF3D7_0811600, which had high ERC and previous interactome-predicted interactions with RhopH2, RhopH3, and RAP1 [8] (Fig 5).

**Fig 5.**
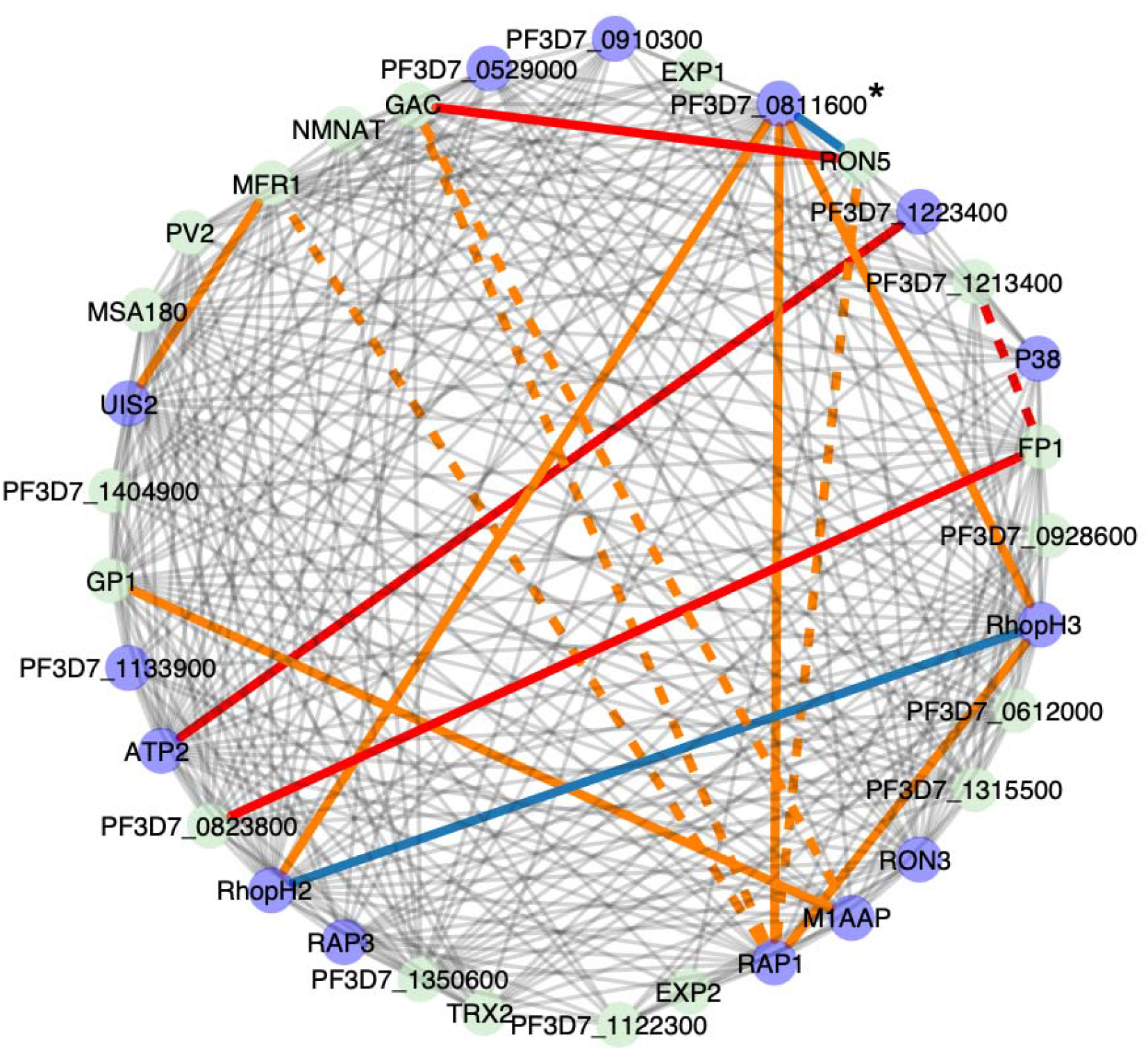
Functional enrichment in cluster 538 of 33 genes with high ERC. Purple nodes have enriched functional annotations. Red lines indicate STRING interactions as described in Fig 3. Orange lines indicate protein pairs that also interact in the experimentally derived interactome Solid orange lines between six pairs denote 99.9th percentile of ERC values and dashed lines between four pairs denote 99-95th percentile of ERC values (4.7 - 2.2). Blue lines indicate pairs with all three: 99.9th percentile ERC, interactome, and STRING interactions. The protein that we followed up experimentally is marked with an asterisk.

To functionally validate the predicted interactions between PFD7_0811600 and RhopH2/RhopH3 suggested by high ERC, we obtained a *P. falciparum* line with an endogenously modified PF3D7_0811600 gene to encode a triple hemagglutinin (3xHA) tag and a glmS ribozyme sequence in the 3’ untranslated region, allowing inducible knock down of gene expression upon addition of glucosamine [49]. Addition of glucosamine to tightly synchronized ring-stage parasites reduced protein levels by 80% within the same cycle and remained constant over multiple life cycles (Fig 6A, B). Parasites showed only a minor growth defect, despite the protein being predicted to be essential in *P. falciparum* and *P. knowlesi* (Fig 6C) [4, 6]. We used immunofluorescence to test for subcellular localization of PFD7_0811600 in fixed parasites and saw an accumulation of the protein at the parasite periphery and in a punctate pattern in the infected RBC in early-stage parasites (Fig 6D). In late-stage parasites, the protein appeared to localize to distinct foci in the forming merozoite. The protein co-localized with both RhopH2 and RhopH3 (Pearson coefficient of 0.6+/-0.0833 and 0.541+/-0.119, respectively), with a very high M1 value (0.93+/-0.0737 with RhopH2 and 0.898+/-0.137 with RhopH3), and a lower M2 value (0.704+/-0.159 with RhopH2 and 0.626+/-0.171 with RhopH3; Fig 6E and Fig S9A). This pattern suggests that PF3D7_0811600 almost entirely localizes proximal to PSAC, but also that there is a subset of PSAC without PF3D7_0811600. We were unable to detect enrichment of RhopH3 in co-immunoprecipitations followed by western blot using PF3D7_0811600 as bait (Fig S9B). This observation, together with the skewed M1 and M2 values, suggests a transient interaction between PF3D7_0811600 and the PSAC.

**Fig 6.**
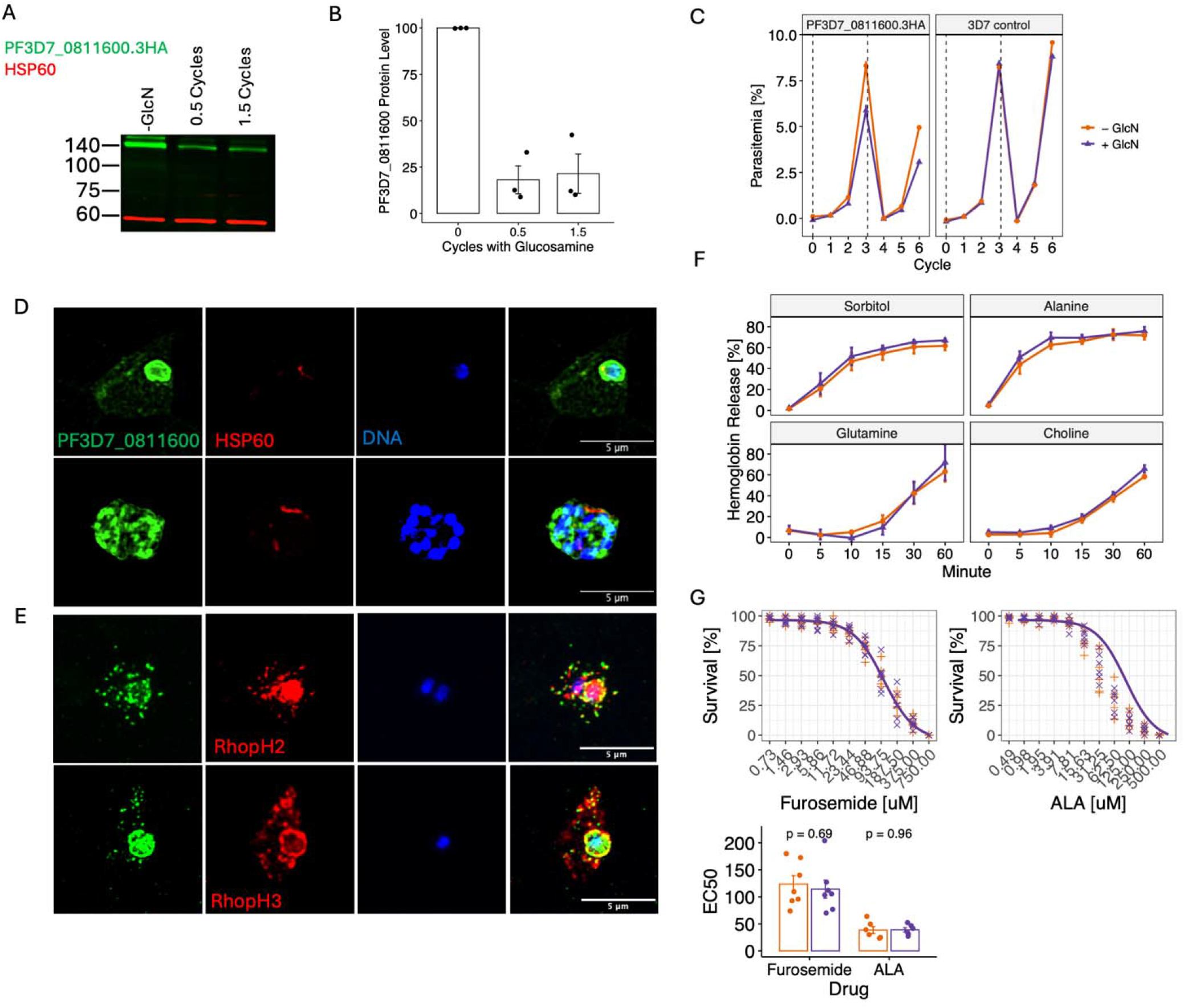
PFD7_0811600 colocalizes with PSAC but does not affect solute import. (A) Western blot of PF3D7_0811600 shows protein knock-down after addition of glucosamine (GlcN) for 0.5 and 1.5 cycles. HSP60 is a marker for the mitochondrion and shown as a control. (B) Quantification of PF3D7_0811600 protein level after knock-down. (C) Growth of parasites with (+ GlcN) and without (-GlcN) knock-down of PF3D7_0811600 and in 3D7 parasites without tagged PF3D7_0811600 (right). (D) Localization of PF3D7_0811600 in infected red blood cells in early-stage parasites (top row) and late-stage parasites (bottom row). (E) Co-localization of PF3D7_0811600 with RhopH2 (top row) and RhopH3 (bottom row). (F) Cell lysis resulting from treatment with solutes, with and without PF3D7_0811600 knock-down. (G) Survival with furosemide and 5-aminolevulinic acid treatment with and without PF3D7_0811600 knock-down.

To test for the role of PF3D7_0811600 in PSAC function, we measured solute import into the infected red blood cell upon protein knock-down. Import of sugars such as sorbitol or amino acids such as alanine require PSAC activity and result in lysis of the cell at high concentrations [50]. Indeed, knock-down of RhopH2 has previously been shown to reduce the uptake of sorbitol and alanine [29]. However, we observed no difference in cell lysis upon PF3D7_0811600 knock-down, quantified as percent of hemoglobin release, after treatment with sorbitol, choline, alanine or glutamine (Fig 6F). This observation mirrors a recent publication where knock-down of none of 13 putative RhopH2 interacting proteins changed sorbitol uptake [51]. We further tested whether knock-down of PFD7_0811600 changes parasite susceptibility to treatment with the PSAC inhibitor furosemide [52] or 5-aminolevulinic acid, which is imported through the PSAC [53], but we did not detect a shift in EC_50_ for either drug (Fig 6G). This observation, together with a lack of change in solute import, suggests that PFD7_0811600 is not directly involved in or required for PSAC function, consistent also with the lack of in vitro growth phenotype upon knock-down. Alternatively, as we only achieved partial knock-down with 20% of the protein remaining within the cell (Fig 6A, B), these low amounts of protein may be sufficient to facilitate its function.

## Discussion

We have generated a pairwise ERC dataset for 4,360 *Plasmodium* proteins calculated across 22 *Plasmodium* species. We found that diverse protein pathways across multiple lifecycle stages show signatures of high ERC, demonstrating this evolutionary-based approach to be a useful screening and/or discovery tool for *Plasmodium* protein function, as in other eukaryotic taxa [17-23]. We illustrated this utility of ERC as a screening tool by two approaches. First, we scanned genome-wide for proteins showing high ERC with co-functional proteins involved in RBC invasion to identify a set of new candidates, which are enriched for blood stage expression and physical interactions. Second, by clustering the highest ERC values genome-wide, we identified uncharacterized proteins that cluster with established co-functional proteins. From this, ERC predicted interaction between PSAC components RhopH2/H3 and a recently characterized protein, PF3D7_0811600, which we confirmed to colocalize.

Recent work found PF3D7_0811600 localized to the parasitophorous vacuole membrane (PVM), which separates the parasite from the host cell cytosol, and knock-down of PF3D7_0811600 led to reduced CD36-mediated cytoadhesion [49]. Similarly, the *P. berghei* ortholog of PF3D7_0811600, called Merozoite Attachment Protein 1 (MAP1), also localized to the PVM and loss resulted in reduced sequestration. PF3D7_0811600 is also referred to as PfMAP1 and is annotated as “protein MAP1, putative” in PlasmoDB. Here we use gene names from PlasmoDB release 54 annotations where it is unnamed. The colocalization of some but not all PSAC components with PF3D7_0811600 suggests a transient or heterogeneous interaction between these proteins. For example, PF3D7_0811600 could be involved in trafficking of PSAC to the host RBC membrane and dissociate after PSAC installation on the RBC surface. To reach the host RBC membrane, RhopH2 and RhopH3 pass through the *Plasmodium* translocon of exported proteins (PTEX) [26], which is a protein complex that transports proteins across the PVM and into the host cytosol [54]. In two previous studies, PF3D7_0811600 was among the proteins that immunoprecipitated with a PTEX intermediate complex, suggesting it could interact with the PTEX components [55, 56]. Twelve of the other 29 immunoprecipitated proteins identified in Miyazaki et al. [55] had ERC values in the 99th percentile (ERC > 3.2) with PF3D7_0811600 (Table S12, Table S13). These proteins included both RhopH2 and RhopH3, HSP101 and EXP2 (two of three core PTEX components) [57], RON3 which is essential for PTEX protein transport function [58], EXP1 which is required for correct localization of core PTEX component EXP2 [59], and PTEX associated protein P113. Seven of the 19 immunoprecipitated proteins identified in Mesén-Ramirez et al [56] had ERC in the 99th percentile with PF3D7_0811600, again including HSP101 and EXP2, and PTEX88, an accessory PTEX component (Table S14). Interestingly, neither study pulled out the second accessory PTEX component, TRX2, yet we observe strong ERC between TRX2 and other PTEX components. However, we do not see strong ERC between core PTEX protein PTEX150 and the other four PTEX proteins, despite its function as a physical adapter linking HSP101 and EXP2 [60] (Fig S10). Further, GO enrichment of proteins in the 99th percentile of ERC with PF3D7_0811600 (Table S12) revealed significant terms including establishment of localization, organic substance transport, transmembrane transport, and host cell cytoplasm (Fig S11, Table S15, Table S16). While we did not see an effect of PF3D7_0811600 on transport of solutes tested in this study (sorbitol, choline, alanine, glutamine, 5-aminolevulinic acid), it is possible that PF3D7_0811600 is involved in transport of other solutes, as the GO enrichment for transporter functions suggests, or that sufficient activity remained even after knocked down and a full knockout would be required to observe an effect on transport. Alternatively, PF3D7_0811600 could be more generally involved in global export of proteins as high ERC with PTEX components and immunoprecipitation with intermediate PTEX complexes suggest.

High ERC can reflect both direct and indirect protein-protein interactions. Here, we observed high ERC between direct, physically binding protein-protein interactions, for example: p12 and p41 (3.7; p = 0.005) [61] and the RON complex consisting of RON2, RON4, and RON5 (5.1; p = 0.001) [32, 62, 63]. We also saw high ERC between indirect protein interactions such as proteins within the same pathway (Table 1) and co-functional proteins (Fig 4, 5). Previous work in yeast showed that ERC was stronger between indirect protein interactions compared to direct interactions, suggesting nonphysical forces such as changes in evolutionary constraint, expression, codon adaptation, or essentiality are primary drivers of ERC [16].

The high ERC observed between RhopH2 and RhopH3, driven by correlated RERs across the species tree, appears to be loosely correlated with host species. The RhopH2/H3 proteins had accelerated RERs in rodent *(P. berghe*i, *P. vinckei, P. chabaudi, P. yoelli*) and avian malaria parasites (*P. relictum, P. gallinaceum*) and decelerated RERs in the ape-infecting (*P. ovale, P. malariae*, and Laverania clade containing *P. falciparum)* and monkey-infecting species (*P. knowlesi*, and *P. vivax* and *P*.*vivax-like)* (Fig 1A). The core structure of the PSAC, including the RhopH2/H3 complex, is thought to be conserved across *Plasmodium* species [48], but differences in affinities of PSAC inhibitors between *Plasmodium* species suggest it may serve different functions [64, 65]. It is plausible that shifts in RERs of RhopH2/H3, and other highly correlated transport proteins (Fig 2B, Fig 5), could be due to different nutrient availability or requirements imparted by unique host species environments, resulting in alteration of nutrient acquisition mechanisms. Indeed, parasite species differ in sensitivity to drugs targeting metabolic pathways [66], essentiality of transporters [67] and metabolic pathways[68], and complete loss of metabolic pathways [69].

We did not observe elevated ERC for the mismatch repair pathway (Table 1). Although mismatch repair proteins are generally conserved across eukaryotes and show elevated ERC in yeast and mammals [23, 70], the mismatch repair pathway appears to be less conserved in *Plasmodium* [71-74] and may also be under selection related to drug treatment [75]. It is possible that our mismatch repair protein list does not show elevated ERC because it contains proteins that serve other functions in *Plasmodium*. Within our mismatch repair protein list, there are subsets of proteins that have high ERC. For example, a group of putative exonucleases (PF3D7_0204600, PF3D7_1362500, PF3D7_0105900, PF3D7_0204800) and the DNA helicase UvrD (PF3D7_0514100) cluster together with higher ERC values (Fig S6). Additionally, replication proteins (RPA1; PF3D7_0409600, PF3D7_0904800 and RPA3; PF3D7_1442100) cluster together. Some proteins in the heatmap show little ERC with others, including two putative mismatch repair proteins (PMS1; PF3D7_0726300, PF3D7_0505500; MSH6). Proteins within the top 100 ERC values with these two putative mismatch repair proteins were enriched for GO terms related to RNA metabolism and processing. High ERC with PMS1 included three U3 small nucleolar RNA associated putative proteins (UTP7; PF3D7_0722600, UTP12; PF3D7_1448000, UTP4; PF3D7_1333600) which are part of the small subunit processome for ribosome biogenesis (Table S17). These results indicate potential divergent functions of putative mismatch repair proteins in *Plasmodium*. Since ERC is estimated using *Plasmodium* sequences, it is a *Plasmodium*-specific inference of protein function and may be particularly useful to predict the function of divergent proteins in *Plasmodium*.

## Materials and Methods

### *Plasmodium* reference genomes

Amino acid FASTA files (ending in “_AnnotatedProteins.fasta”) were downloaded from PlasmoDB release 50 (https://plasmodb.org/plasmo/app/downloads) for each of 22 *Plasmodium* species: *P. adleri* G01, *P. reichenowi* CDC, *P. praefalciparum* G01, *P. relictum* SGS1, *P. vinckei vinckei vinckei, P. vivax-like* Pvl01, *P. vivax* P01, *P. yoelii yoelii* 17X, *P. ovale curtisi* GH01, *P. malariae* UG01, *P. knowlesi* H, *P. inui* SanAntonio1, *P. gallinaceum* 8A, *P. gaboni* G01, *P. fragile* Nilgiri, *P. berghei* ANKA, *P. billcollinsi* G01, *P. blacklocki* G01, *P. chabaudi chabaudi, P. coatneyi* Hackeri, *P. cynomolgi* M, and *P. falciparum* 3D7 (Table S1). We used the “PlasmoDB-54_Pfalciparum3D7.gff” file to annotate protein names and descriptions also downloaded from the link above.

### Ortholog assignment and alignment

Ortholog groups within the 22 species were assigned using OrthoMCL v2.0.91 [76]. Protein sequences flagged as short or highly repetitive by OrthoMCL were excluded. For proteins with multiple isoforms, a single isoform was kept (selecting the isoform annotated with ‘.1’). Paralogs were removed and only ortholog groups with at least ten 1:1 orthologs were kept. We refer to ortholog groups by the *P. falciparum* gene name when one is present. If an ortholog group did not contain a single-copy *P. falciparum* ortholog, it was named by the *P. berghei* ortholog, followed by *P*.*vivax, P. reichenowi, P. coatneyi, P. malariae, P. ovale, P. knowlesi, P. cynomolgi, P. inui, P. fragile, P. chabaudi, P. yoelii, P. vinckei, P. gaboni, P. adleri, P. blacklocki, P. billcollinsi, P. praefalciparum, P. vivax-like, P. relictum, P. gallinaceum* ortholog. Protein sequences of each one-to-one ortholog group were aligned using MUSCLE v3.8.31[77].

### Relative evolutionary rate and evolutionary rate covariation estimates

The species topology was inferred using published phylogenetic trees available for subsets of species as well as literature consensus [78-81] (Fig 1A). This species topology and the multiple sequence alignment of each ortholog group were used to estimate individual maximum likelihood gene trees with the estimatePhangornTreeAll wrapper within the R package RERconverge (v0.1.0) [10, 82], with default options and the LG amino acid substitution matrix [83]. Protein relative evolutionary rates (RERs) were estimated by normalizing a gene’s branch length in each species to its average evolutionary rate across the species tree and to the average rate of all other genes on that branch, detailed elsewhere [10, 11, 84, 85]. Evolutionary rate covariation (ERC) was calculated as the Pearson correlation of the RERs between two ortholog groups across the phylogeny, followed by a Fisher transformation to normalize for comparisons between ortholog groups with different numbers of branches, as previously described [16, 22]. ERC was calculated between all pairs of ortholog groups with at least 10 shared species. The resulting genome-wide matrix of protein-protein pairwise ERC values was used in all downstream analyses and can be found under data availability.

### ERC analysis of proteins in known pathways

Lists of proteins with established roles in pathways were obtained from Malaria Parasite Metabolic Pathways (MPMP) databases [34] (Table S2). Specifically, under “maps” and “map details”, we extracted the gene names under fatty acid synthesis II (https://mpmp.huji.ac.il/maps/facidsynthesispath.html), DNA replication (https://mpmp.huji.ac.il/maps/dnareplication.html), double strand break repair and homologous recombination (https://mpmp.huji.ac.il/maps/dsbRecomb.html), hemoglobin digestion (https://mpmp.huji.ac.il/maps/hemoglobinpolpath.html), and DNA mismatch repair (https://mpmp.huji.ac.il/maps/DNArepair.html) pathways. The red blood cell invasion gene set was compiled from the MPMP website (https://mpmp.huji.ac.il/maps/Merozoiteproteins.html) and several review papers [28, 32, 33]. This set includes both proteins with both known functions in red blood cell invasion and those with putative roles due to their localization to rhoptries, micronemes, or the merozoite surface (Table S2). To assess the strength of ERC for each pathway, we calculated the mean ERC across all pairs of pathway proteins. We then randomly sampled gene sets of the same size and computed the pairwise mean ERC in the same way, repeating 1000 times to determine an empirical p-value. This same process was repeated for *P. falciparum* STRING network clusters downloaded from https://string-db.org/.

### Comparison with an experimental interactome

We downloaded a *Plasmodium* protein interactome derived from experimental protein co-migration combined with other evidence for interaction including co-expression, interacting domains, co-evolution, and phenotype data [8]. To assess the overlap between high ERC values and interactions identified in the interactome, we only considered protein pairs that were included in both datasets. A majority (2226/2672, 83%) of the *P. falciparum* proteins queried in the interactome dataset were also in the ERC matrix. About half (2226/4134, 53%) of the *P. falciparum* proteins in the ERC matrix were also queried in the interactome dataset. The mean ERC value of the 17,174 *P. falciparum* protein interactions from the interactome dataset was calculated and an empirical p-value determined similarly as for pathways, except that proteins were randomly sampled only from those that were also in the interactome.

### Analysis of top ERC values genome-wide and with RBC invasion proteins

We selected the 99.9th percentile of ERC values (> 4.48) across the entire dataset to explore genome-wide patterns of the strongest ERC values. To identify groups of potential co-functional proteins among the top genome-wide ERC values, we applied MCL clustering followed by Leiden clustering to the ERC pairs using clusterMaker2 and visualized the clusters within cytoscape (v3.10.1) [86]. For functional enrichment analyses, we used clusterProfiler (v4.12.6) with the *P. falciparum* annotation database package org.Pf.plasmo.db (v3.19.1) for Gene Ontology (GO) enrichment [87]. We also used the STRING annotation databases and the get_enrichment function in the STRINGdb R package (v2.16.4) for functional enrichment. We used all *P. falciparum* orthologs in the ERC matrix as the background.

We also selected the 99.9th percentile of ERC values with known RBC invasion proteins to evaluate for functional candidates and co-functional relationships in the process of RBC invasion. We tested for an enrichment of pairs also in the interactome pairs by randomly sampling sets of the same number of ERC pairs from the interactome filtered ERC dataset described above. To test for enrichment of expression of RBC invasion ERC candidates in blood stages, we used *P. falciparum* single cell gene expression data from the Malaria Cell Atlas covering the asexual blood stages (rings, schizonts, trophozoites) [88]. We considered a gene expressed in a particular stage if it was expressed in at least 25% of cells from that stage. To intersect our RBC invasion candidates with previously identified candidates based on transcriptional patterns, we downloaded supplementary table 6 from Hu et al. 2010 [35] and converted older gene identifiers using PlasmoDB release 54 gene aliases. The RBC invasion genes used as the well-established reference genes in their paper were excluded prior to intersection with our candidates.

### Parasite culture, transfection, growth assays and drug assays

Parasite culture, transfection and selection were performed as previously described [89, 90]. Parasites were cultured in RPMI1640 supplemented with 2.5 g/L Albumax I Lipid-Rich BSA, 15 mg/L hypoxanthine, 110 mg/L sodium pyruvate, 1.19 g/L HEPES, 2.52 g/L sodium bicarbonate, 2 g/L glucose, and 10 mg/L gentamicin. Cultures were maintained at 2.5% hematocrit in human erythrocytes obtained from the University of Utah Hospital blood bank, at 37°C, and 5% CO2. Parasites were kept synchronous through treatment with 5% (w/v) Sorbitol. Parasite growth and viability was routinely controlled via Giemsa stained thin blood smear.

Knock-down of PF3D7_0811600 was initiated by addition of 2.5mM glucosamine [91]. Late-stage PF3D7_0811600.3HA.glms parasites were enriched using MACS magnetic columns [92]. Continuous growth assays and drug assays were performed as previously described [93].

Parasites treated with ALA were subjected to 2 exposures to broad-wavelength white light (2 minutes each, 24 and 36 hours after addition of ALA) on an overhead projector [53].

### Western blot analysis

Late-stage parasites were lysed in PBS with 1% Triton X-100, 0.1% Sodium dodecyl sulfate, 1:1000 Pierce Universal Nuclease for Cell Lysis and 1:200 SelleckChem Protease Inhibitor Cocktail (EDTA free, 100X in DMSO) for 30 minutes at 4C. Lysates were clarified through centrifugation at 13.000g for 30 minutes at 4C. Clarified lysates were mixed with loading dye loaded onto 10% polyacrylamide gels and fractionated via SDS-PAGE. Proteins were transferred onto nitrocellulose membrane. Membranes were airdried, rehydrated in PBS and then blocked in PBS with 5% skimmed milk powder for 30 minutes at room temperature. Membranes were incubated with primary antibodies diluted in PBS + 1% skimmed milk + 0.1% Tween20 overnight at 4C with gentle agitation. Membranes were washed thrice with PBS + 0.1% Tween20 and incubated with secondary antibodies for 1hr at room temperature. Secondary antibodies were diluted in PBS + 1% skimmed milk + 0.1% Tween20. Membranes were washed again thrice with PBS + 0.1% Tween20, rinsed with PBS and airdried. Membranes were scanned on a Li-Cor Odyssey CLx. Band intensities were measured using Li-Cor Image Studio Version 5.2. To pre-clear primary antibody solutions containing antibodies against RhopH2 or RhopH3, samples of uRBCs were prepared and processed as described above. Primary antibody solutions were incubated with membranes loaded with uRBC samples for 1hr at room temperature. Pre-cleared primary antibody solutions were then transferred onto membranes with the samples of interest.

### Immunofluorescence analysis

All steps were performed at room temperature. Parasite cultures were washed once using PBS and diluted to 0.1% hematocrit in PBS. Parasite suspension was transferred to Poly-□ - Lysine Hydrobromide (molecular weight 30,000-70,000) coated cover slips and left to settle for 30 minutes. Coverslips were rinsed with PBS and cells were fixed with 4% paraformaldehyde in PBS for 30 minutes. Fixed cells were permeabilized with PBS + 0.1% Triton-X100 for 5 minutes. Cells were blocked with PBS + 3% BSA for 30 minutes, followed by incubation with primary antibodies in PBS + 1% BSA for 1hr. Cover slips were washed twice with PBS for 5minutes. Cells were finally incubated with secondary antibodies and Hoechst33342 (1 ug/ml) in PBS + 1% BSA for 1hr. After two more washes in PBS for 5 minutes, cells were sequentially dehydrated in 70%, 90% and 100% ethanol for 3 minutes each. Cover slips were left to airdry and mounted onto glass slides using Invitrogen ProLong Diamond Antifade Mountant with DAPI. Cells were imaged within the next 7 days on a Nikon ECLIPSE Ti2-E inverted microscope with a Yokogawa CSU-W1 Confocal Scanner Unit (with 50um pinhole disk), a Gataca Systems Live-SR Super Resolution Module and an Apo TIRF 100x Oil DIC N2 objective using a Teledyne Kinetix22 Back Thinned sCMOS Camera (6.5um pixel size and 2400×2400 sensor). Images were taken using the Nikon NIS elements AR/HC software package (versions 5.42 or 6.02) and deconvoluted using NIS elements Batch Deconvolution (version 5.30.01) in automatic mode. Images were analyzed using FIJI [94] (Version 2.16.00, Java 1.8.0_322) and colocalization was calculated using the plugin JaCOP [95] (Version 2.1.4).

### Co-immunoprecipitation

To detect direct protein-protein interactions between PF3D7_0811600 and RhopH2 or RhopH3, late-stage parasites were harvested using MACS magnetic columns [92]. Parasites were lysed in PBS with 1% Triton X-100, 1:1000 Pierce Universal Nuclease for Cell Lysis and 1:200 SelleckChem Protease Inhibitor Cocktail (EDTA free, 100X in DMSO) for 30 minutes at 4C. Lysates were clarified through centrifugation at 13.000g for 30 minutes at 4C. Clarified lysates were mixed with Pierce Anti-HA Magnetic Beads at a 1:10 ratio (1 volume bead slurry: 10 volumes infected RBC pellet) and incubated at 4C overnight with gentle agitation. Beads were washed 5 times with ice cold PBS with 1% Triton X-100 for 5 minutes at 4C with gentle agitation. Bound proteins were eluted in non-reducing loading dye for 5 minutes at 95C. Samples were supplemented with 2-mercaptoethanol to a final concentration of 3% (v/v) and incubated at 95C for 5 minutes. Western blotting was performed as described above.

### Hemoglobin release assay

The hemoglobin release assay to assess PSAC function was performed as previously described [52]. In brief, late-stage PF3D7_0811600.3HA.glms parasites (untreated or treated with glucosamine for 48 hours) were harvested using MACS magnetic columns [92]. 1/7^th^ of the parasite pellet was incubated in PBS with 1% Triton X-100 for 15 minutes at room temperature to achieve 100% hemoglobin release. The remaining parasites were incubated with either D-sorbitol (300mM), L-alanine (300mM), L-glutamine (300mM) or choline chloride (160mM) at 37C. Aliquots of cell suspensions were taken at 0, 5, 10, 15, 30 or 60 minutes. Cells and debris were pellet for 30 seconds at 10.000 x g and supernatant was transferred to 96 Well flat bottom assay plates. Absorbance at 540nm was measured using a BioTek Synergy Neo2 plate reader.

Hemoglobin release is given as fraction of maximum release (100%, defined as the release from treatment with 1% Triton X-100).

## Acknowledgements

The support and resources from the Center for High Performance Computing at the University of Utah are gratefully acknowledged. We thank Ellen SC Bushell and Julian C Rayner for providing parasite lines as well as comments and suggestions.

## Data availability

Dryad: https://doi.org/10.5061/dryad.cfxpnvxmw

