## Supplemental Figures 1-11 for "Evolutionary rate covariation across malaria parasite species enables inference of protein interactions"

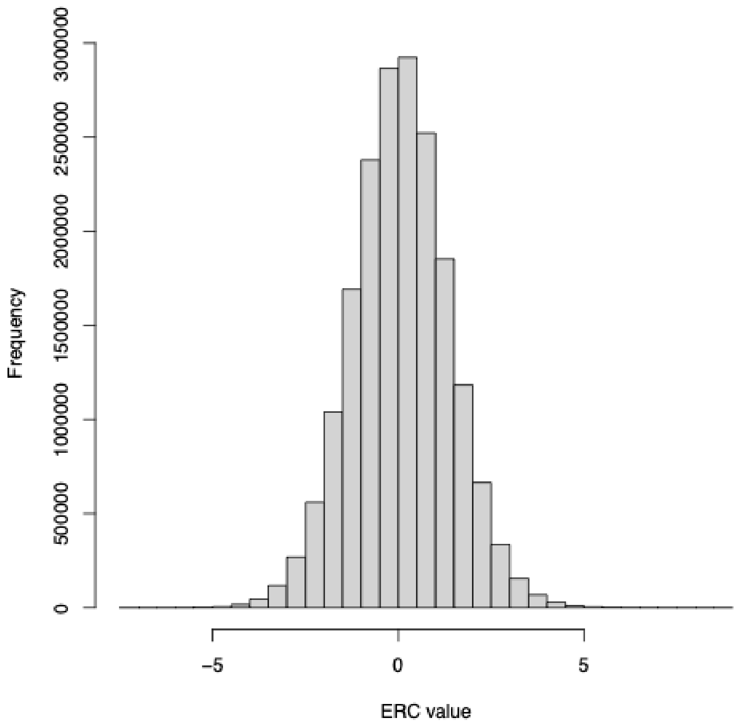


Fig S1. Distribution of ERC values in the whole matrix.

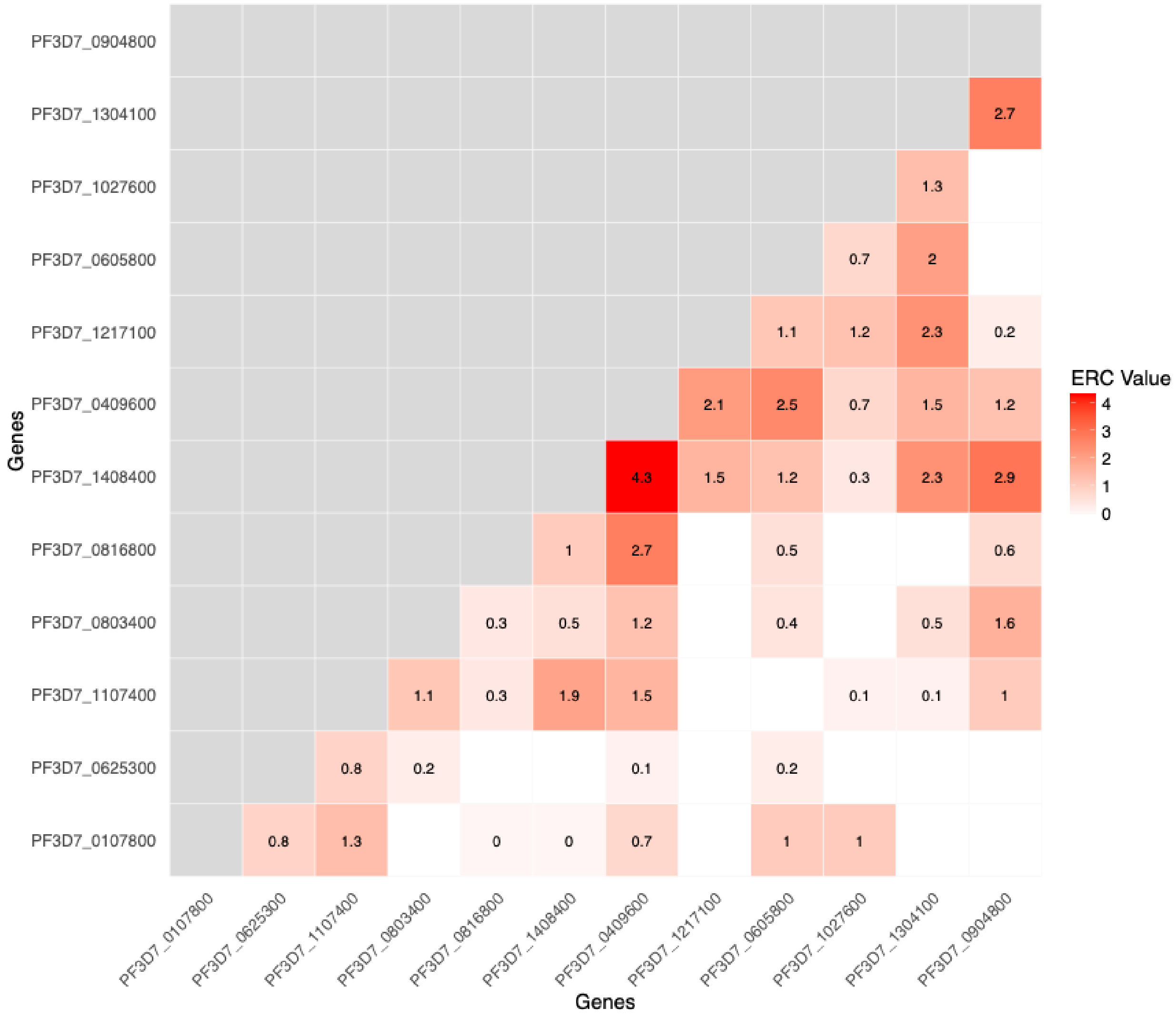


Fig S2. Pairwise ERC between components of the homologous recombination pathway. White squares indicate negative ERC values.

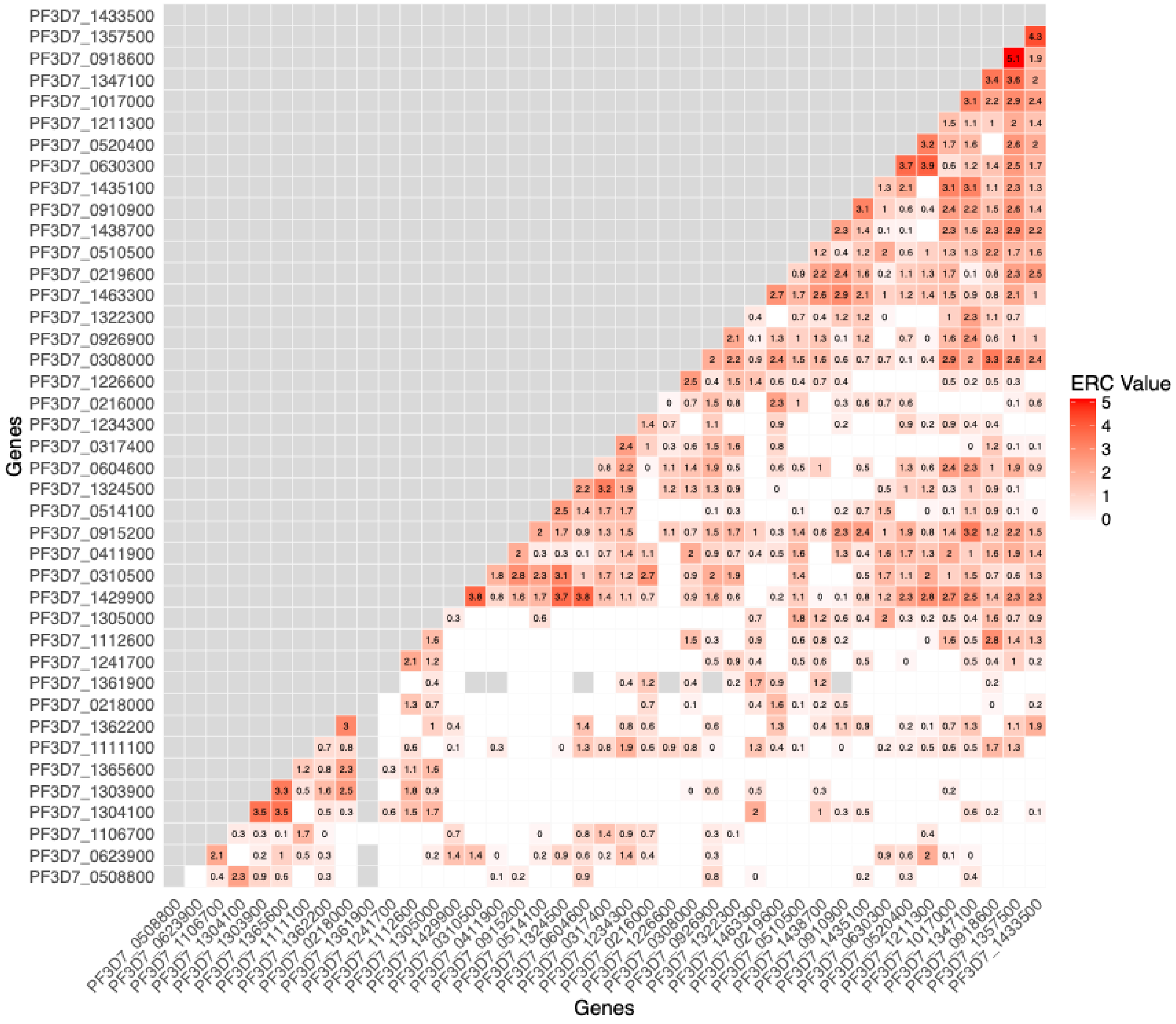


Fig S3. Pairwise ERC between proteins in the DNA replication pathway. Grey squares indicate ERC values were not calculated (proteins did not share 10 species with 1-to-1 orthologs) and white squares indicate negative ERC values.

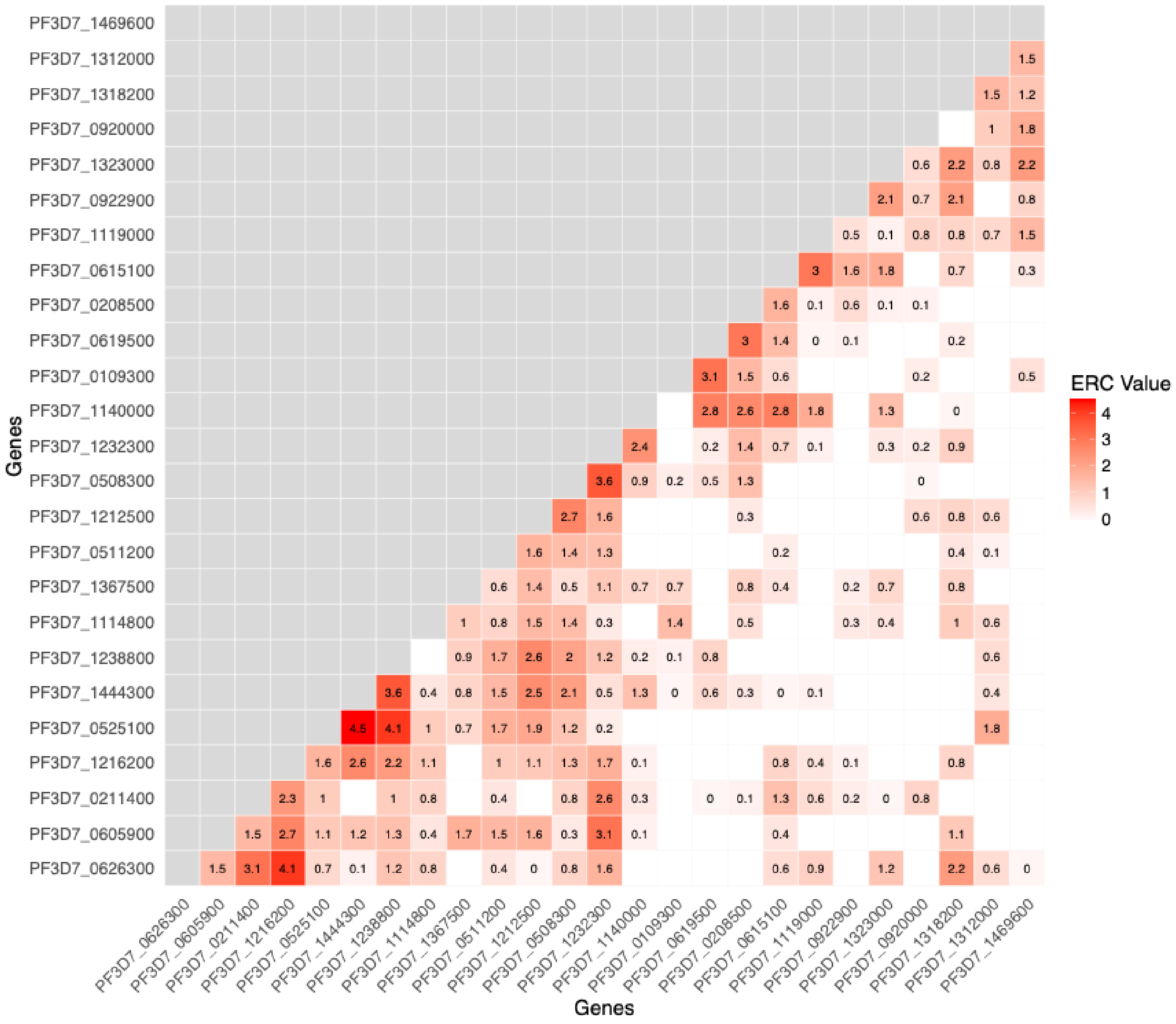


Fig S4. Pairwise ERC between proteins in the fatty acid synthesis II pathway. White squares indicate negative ERC values.

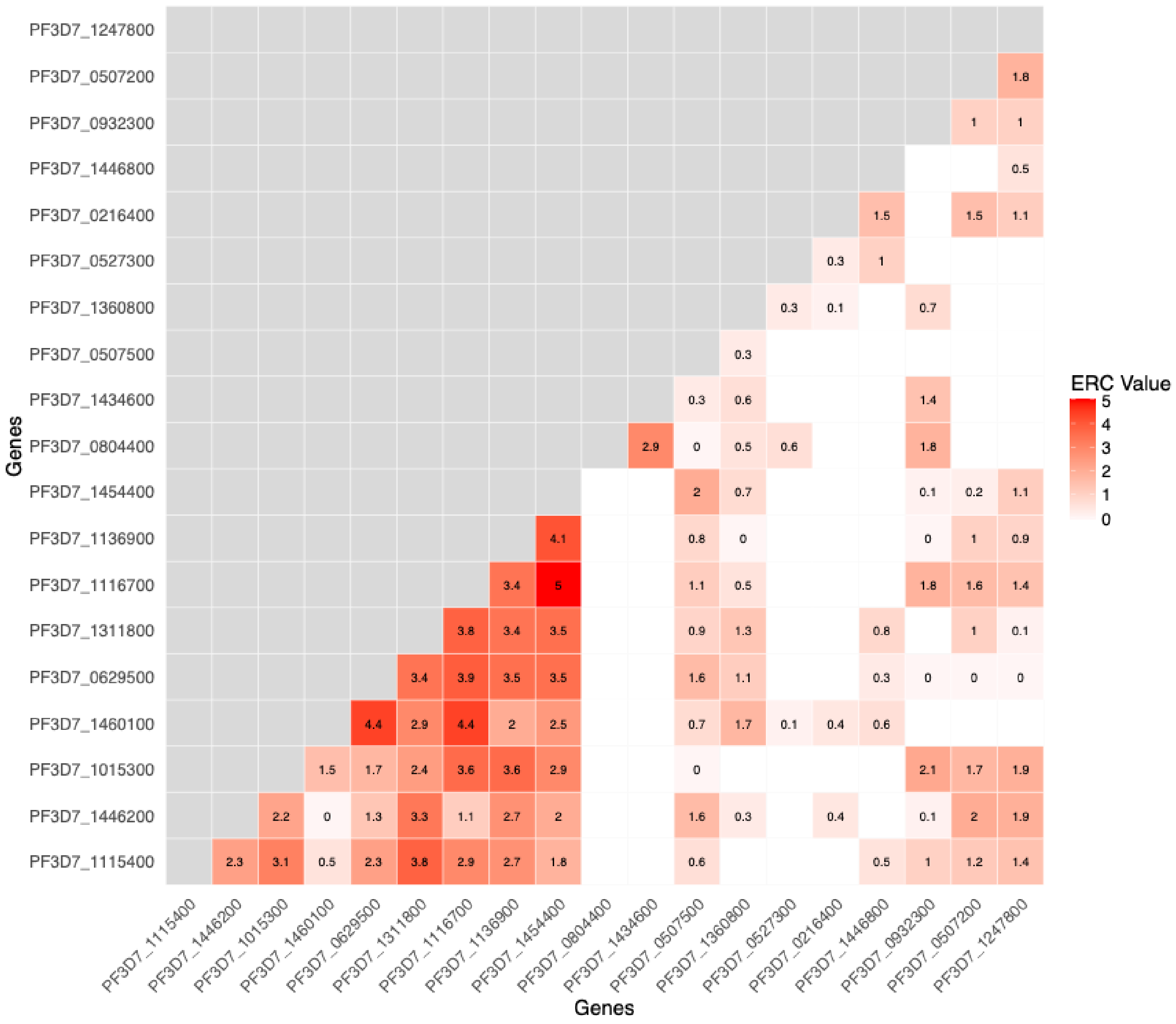


Fig S5. Pairwise ERC between proteins in the hemoglobin digestion pathway. White squares indicate negative ERC values.

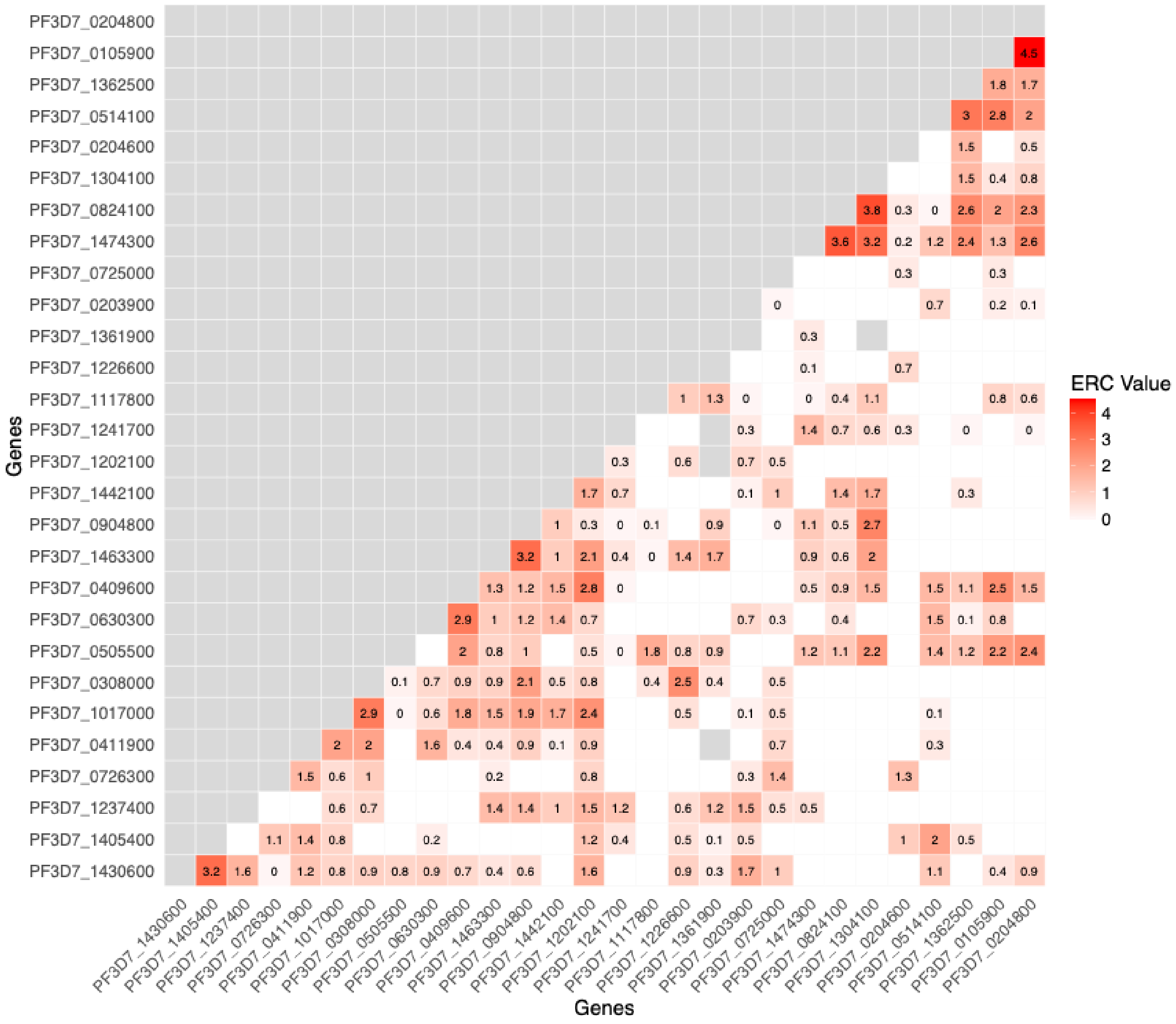


Fig S6. Pairwise ERC between proteins in the mismatch repair pathway. Grey squares indicate ERC values were not calculated (proteins did not share 10 species with 1-to-1 orthologs) and white squares indicate negative ERC values.

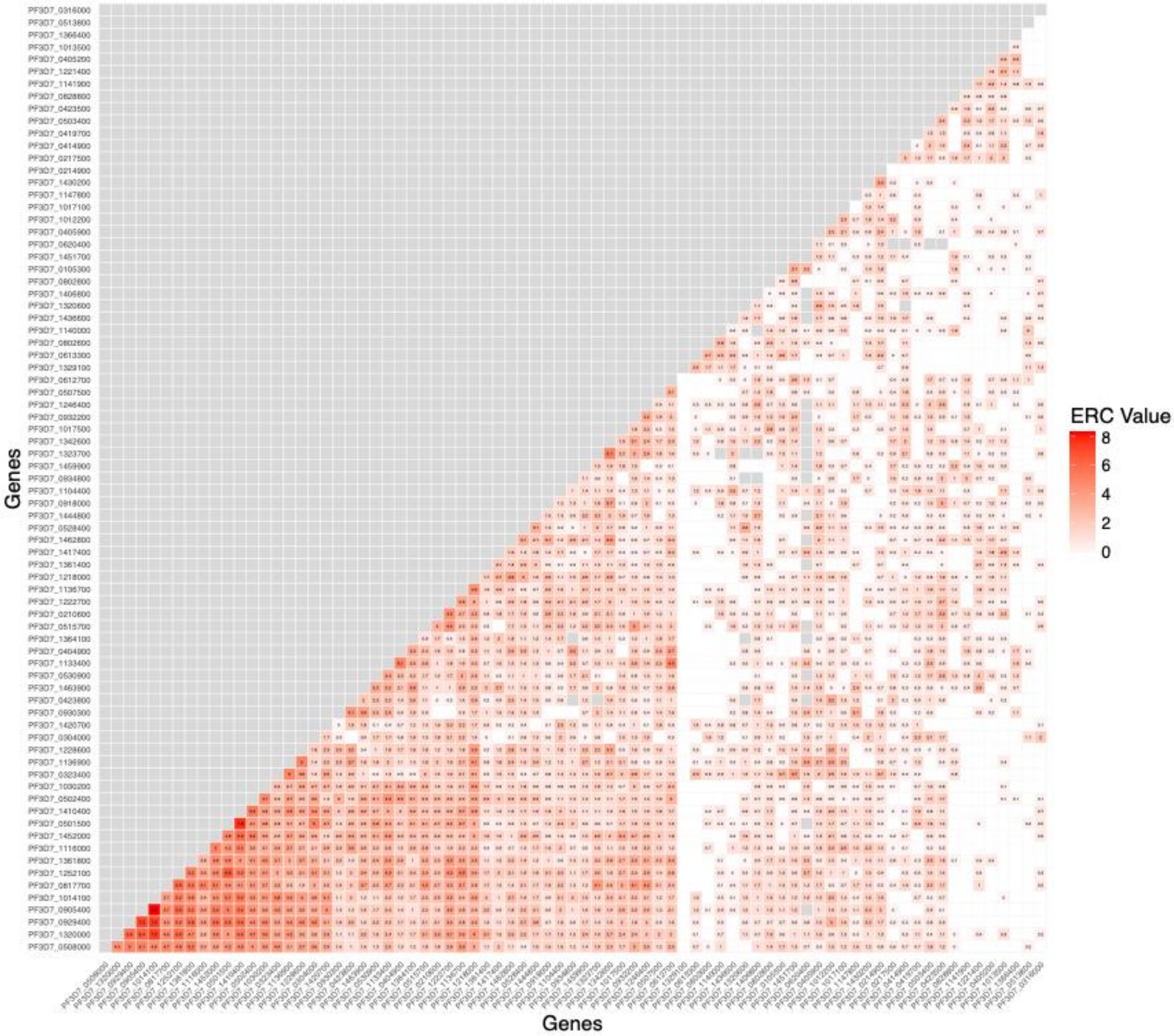


Fig S7. Pairwise ERC between proteins in RBC invasion pathways. Grey squares indicate ERC values were not calculated (proteins did not share 10 species with 1-to-1 orthologs) and white squares indicate negative ERC values.

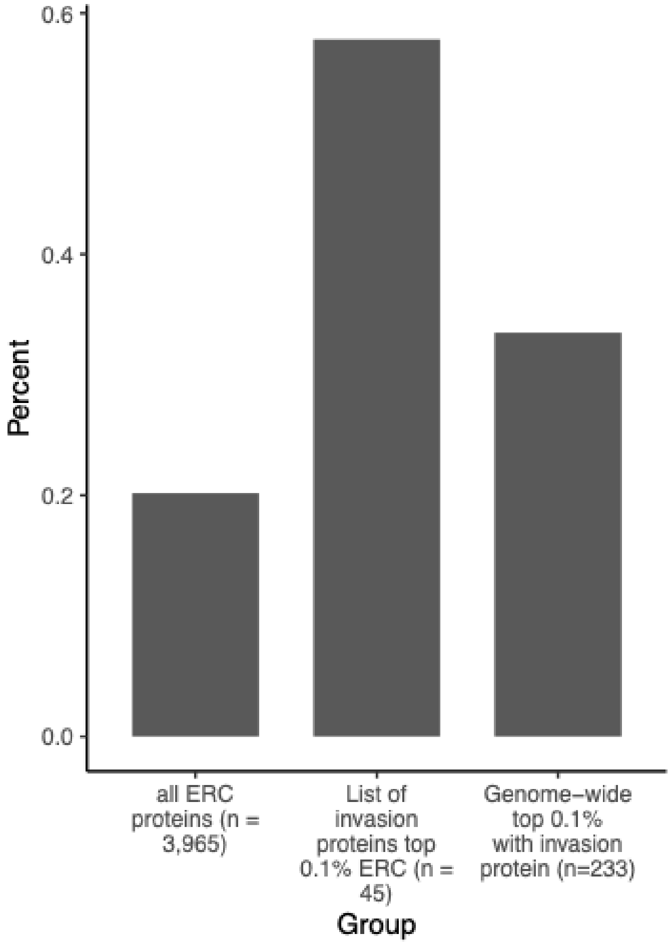


Fig S8. Enrichment of blood-stage expression in RBC invasion protein candidates inferred from ERC. Numbers in each group are after filtering to only keep proteins with single cell expression data.

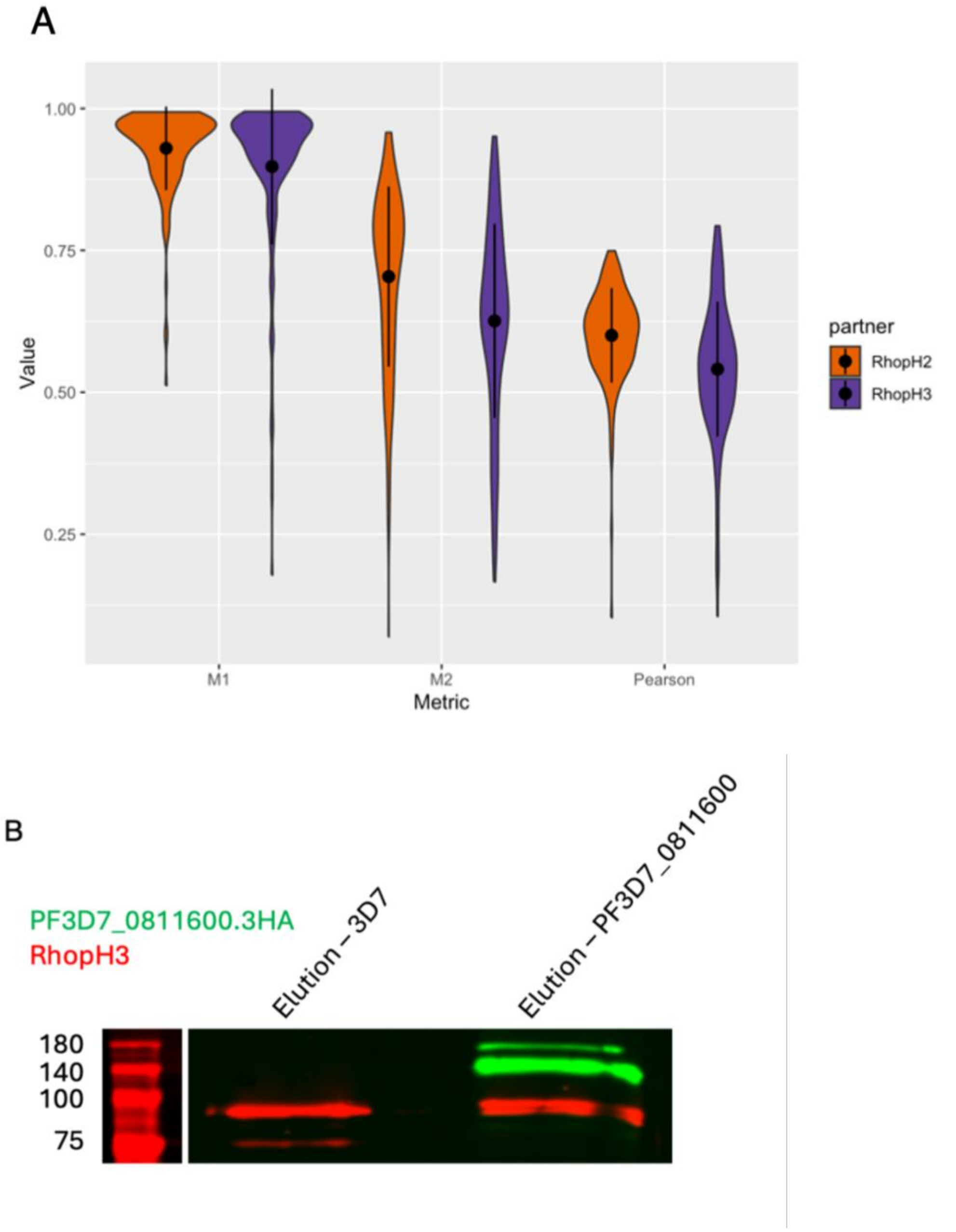


Fig S9. (A) Quantification of co-localization of PF3D7_0811600 with either RhopH2 (orange, n=158) or RhopH3 (purple, n=159) in IFAs. Cells were imaged in three independent replicates. M1 describes how often PF3D7_0811600 co-localizes with either RhopH2 or RhopH3 and M2 describes how often Rhoph2 or RhopH3 co-localize with PF3D7_0811600. (B) Western blot result of a Co-IP, using PF3D7_0811600 as bait and probing for RhopH3 using anti-HA magnetic beads. IP was performed in parental 3D7 and the tagged cell line. No enrichment of RhopH3 could be seen in the tagged cell line compared to the parental control.

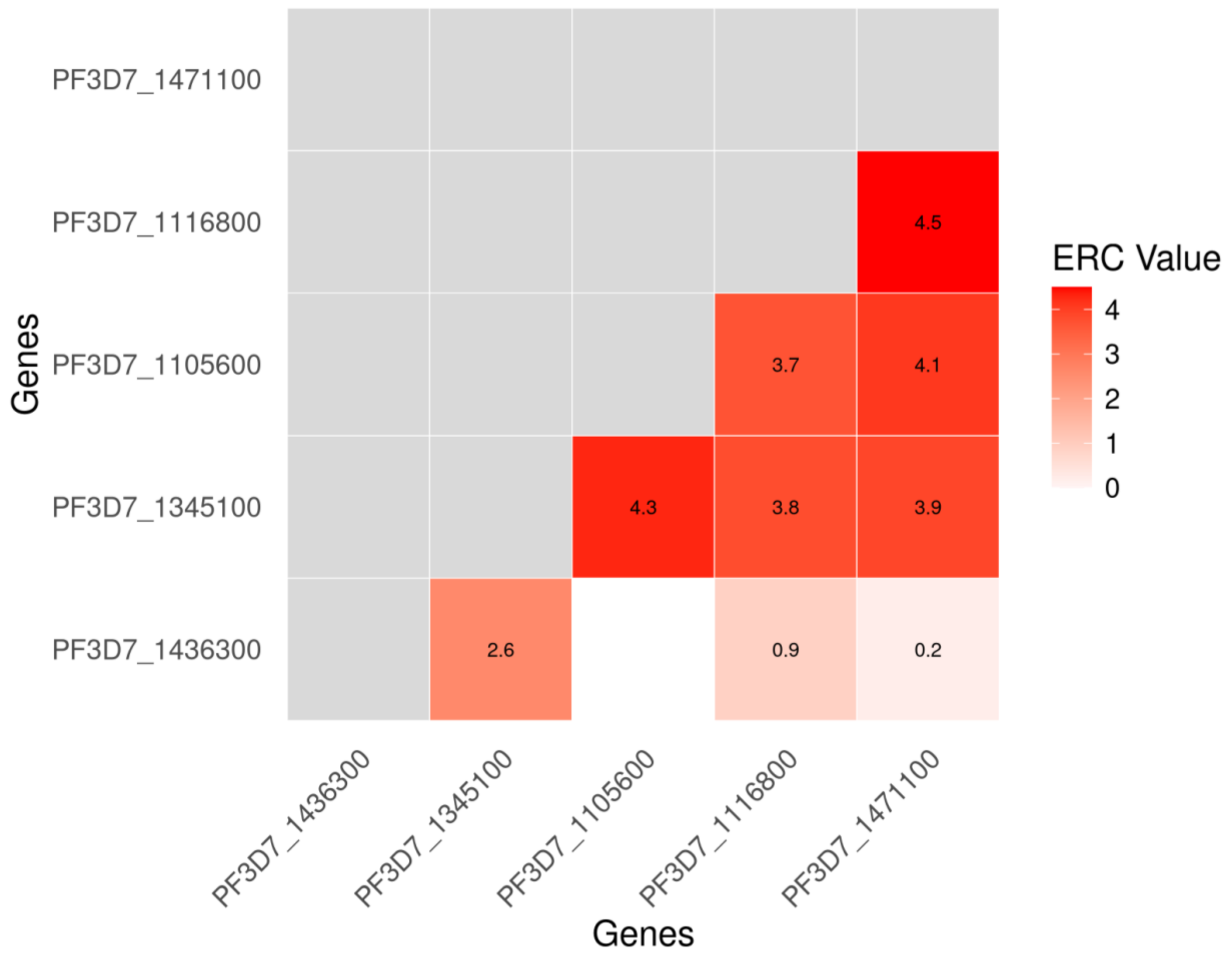


Fig S10. Pairwise ERC between proteins in the PTEX complex. White squares indicate negative ERC values. PTEX proteins listed from left to right: PTEX150, TRX2, PTEX88, HSP101, EXP2.

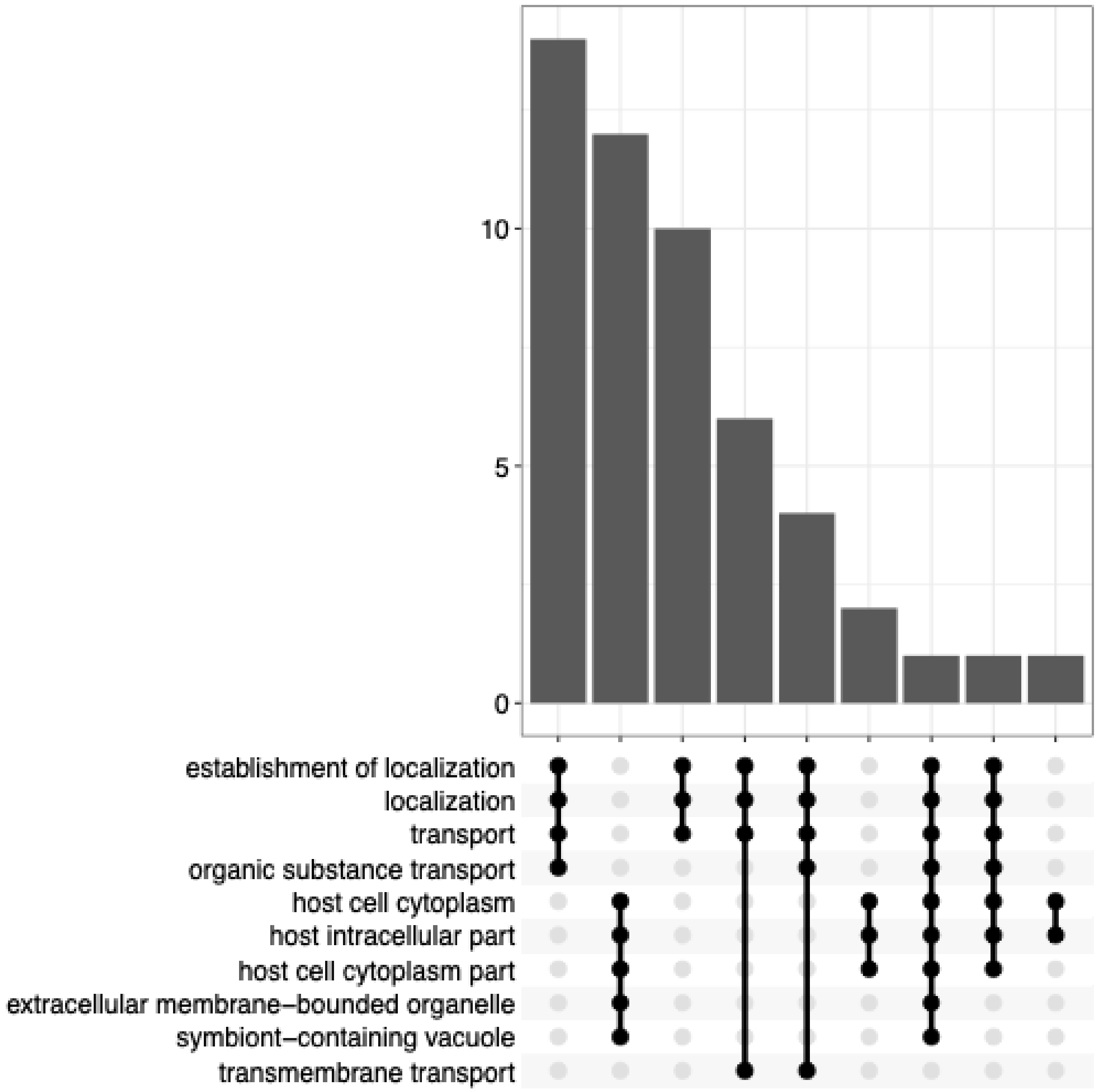


Fig S11. Upset plot of GO enrichment of proteins showing ERC with PF3D7_0811600 99th percentile of ERC values (>3.2).
